# Topographic connectivity and cellular profiling reveal detailed input pathways and functionally distinct cell types in the subthalamic nucleus

**DOI:** 10.1101/2021.10.01.462690

**Authors:** Hyungju Jeon, Hojin Lee, Dae-Hyuk Kwon, Jiwon Kim, Keiko Tanaka-Yamamoto, Linqing Feng, Hyeran Park, Yong Hoon Lim, Zang-Hee Cho, Sun Ha Paek, Jinhyun Kim

**Affiliations:** Center for Functional Connectomics, Korea Institute of Science and Technology (KIST), Seoul, 02792, Korea; Division of Bio-Medical Science & Technology, KIST-School, University of Science and Technology, Seoul, 02792, Korea; Neuroscience Convergence Center, Korea University, Seoul, 02841, Korea; Soonchunhyang University Seoul Hospital, Seoul, 04401, Korea; Neurosurgery, Movement Disorder Center, Seoul National University College of Medicine, Advanced Institute of Convergence Technology (AICT), Seoul National University, Seoul, 03080, Korea

**Keywords:** Subthalamic nucleus, connectivity, indirect and hyperdirect pathways, cell type, firing pattern

## Abstract

The subthalamic nucleus (STN) controls psychomotor activity and is an efficient therapeutic deep brain stimulation target in Parkinson’s disease patients. Despite evidence indicating position-dependent therapeutic effects and distinct functions within the STN, input circuit and cellular profile in the STN remain largely unclear. Using advanced neuroanatomical techniques, we constructed a comprehensive connectivity map of the indirect and hyperdirect pathways in both the mouse and human STN. Our detailed circuit- and cellular-level connectivity revealed a topographically graded organization with three convergent types of indirect and hyperdirect-pathways. Furthermore, we identified two functional types of glutamatergic STN neurons (parvalbumin, PV +/- neurons) segregated with a topographical distribution. Glutamatergic PV+ STN neurons contribute to burst firing. We confirmed synaptic connectivity from indirect and hyperdirect pathways to both PV+ and PV-. These data suggest a complex interplay of information integration within the basal ganglia underlying coordinated movement control and therapeutic effects.

## Introduction

The subthalamic nucleus (STN), a key element of the basal ganglia (BG), plays a central role in psychomotor control and serves as the most efficient therapeutic target of deep brain stimulation (DBS) in Parkinson’s disease patients (Benabid et al., 2009; Bergman et al., 1990; Bevan, 2017; Eisenstein et al., 2014; Group, 2001; Herzog et al., 2004; Kopell et al., 2006; Limousin et al., 1995; Lozano and Lipsman, 2013). Within the STN, two principal tracts modulate the activity of the BG: the indirect and hyperdirect pathways. The indirect pathway carries most GABAergic inputs to the STN from external globus pallidus (GPe) neurons (Mallet et al., 2012; Smith et al., 1998), while the hyperdirect pathway carries most glutamatergic inputs to the STN (**Figure 1A**) (Monakow et al., 1978; Nambu et al., 2002; Smith et al., 1998). The balance of synaptic excitation and inhibition is critical to the operation of brain microcircuits, and this balance is often perturbed in diseases (Chu et al., 2017; Cobos et al., 2005; del Pino et al., 2013; Fan et al., 2012; Kopell et al., 2006; Turrigiano, 2011; Yizhar et al., 2011). The effects of STN DBS appear to involve an interplay of converging inhibitory indirect and excitatory hyperdirect pathways, as well as multiple interactions from molecular and cellular processes that translate into global network effects (Gradinaru et al., 2009; Hashimoto et al., 2003; Hemptinne et al., 2015; Li et al., 2012; Stefani et al., 2005; Windels et al., 2000). Although there is evidence indicating heterogeneous firing patterns, distinct functional movement control, and electrode position-dependent therapeutic effects including side-effects of DBS within the STN (Herzog et al., 2004; Kaku et al., 2019; Kim et al., 2015; Mallet et al., 2012; Mosher et al., 2021; Wodarg et al., 2012), detailed topographical profiling of connectivity, cytoarchitecture, and physiological properties remain incomplete. Thus, comprehensive information regarding these input patterns as well as cellular profiles of the STN is critical for deepening our understanding of normal movement control, Parkinson’s disease, and DBS. Here, using advanced neuroanatomical techniques, we developed detailed three-dimensional (3D) descriptions of the convergent connectivity in the indirect and hyperdirect pathways, as well as molecular and cellular profiles of STN circuitry. Our analysis includes a comparison of ultrahigh-resolution fiber tractography in the human STN with respect to the effectiveness of STN DBS in Parkinson’s disease patients. Our circuit-level connectivity revealed a topographically graded organization, with overlaps and no evident anatomical boundaries, and direct cortico-pallidal connection in both the mouse and human STN. Together with cellular-level connectivity analysis, we found, at least, three convergent types of indirect and hyperdirect-pathway (GPe-only, STN-only, and both) suggesting complex signal integration, such as a classical convergence of inhibition and excitation, feedforward inhibition, and collateral convergence. These circuit features imply functional computational complexity and delicate control. Furthermore, we identified two functional types of glutamatergic STN parvalbumin (PV) +/-neurons that integrate information from both indirect and hyperdirect pathways, are segregated along a topographical distribution. Glutamatergic PV+ neurons show phasic burst firing and are particularly prevalent in the dorsolateral and middle STN, whereas PV- neurons fail to generate burst firing and prevalent in the ventromedial and middle STN. In addition, we confirmed synaptic connectivity from indirect and hyperdirect pathways to PV+/- neurons by combining retrograde virus injections and mammalian GFP reconstitution across synapses (mGRASP) (Kato et al., 2014; Kim et al., 2012). Our analysis corrects errors and adds a critical new level of detail to understanding STN circuitry.

**Figure 1.**
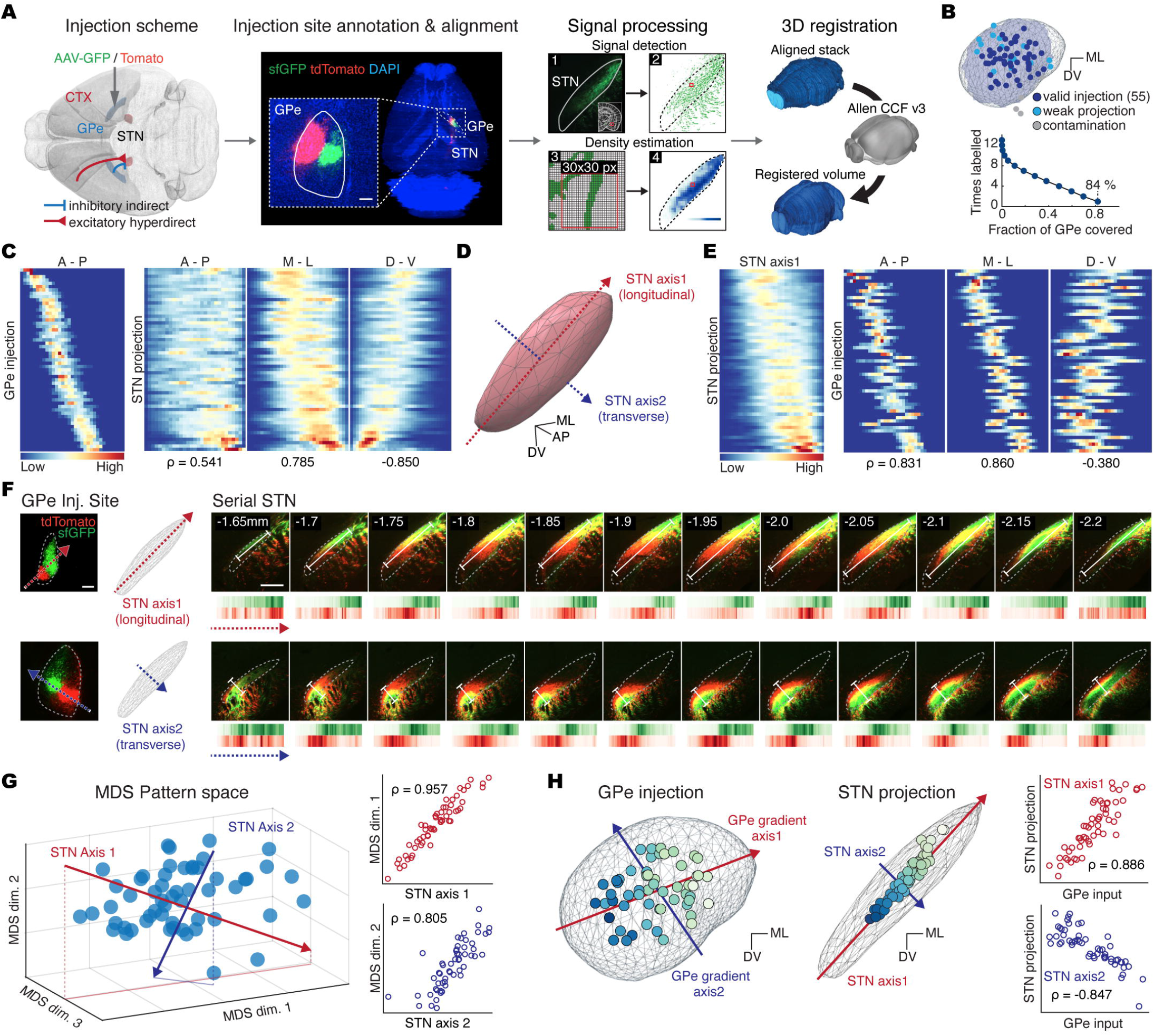
Graded topography of indirect pathway in STN. **(A)** Pipeline of indirect pathway (IP_STN_) mapping data generation and processing. Scale bar, 250 µm. **(B)** Spatial locations (left) and coverage (right) of the GPe injections (n = 55). **(C)** Heatmap showing GPe inputs and IP_STN_ signal distribution for each injection experiment (rows) along anatomical axes (columns), sorted by the GPe injection centroid along the AP. **(D)** Illustration of STN geometric axis1 and axis2. **(E)** Heatmap of IP_STN_ and GPe inputs signal distribution, sorted by projection centroid along STN axis1. **(F)** Representative images of the GPe injection sites (left) and serial coronal images of corresponding IP_STN_ showing graded topographic organization along STN axis1 (right, top) and axis2 (right, bottom). Heatmap showing signal distribution for each slice along the white line, parallel to the corresponding STN axis. The distance from the bregma indicated at the corner. Scale bar, 250 µm. **(G)** Left: IP_STN_ projection patterns (dots) plotted in a pattern space generated via multidimensional scaling (MDS). Right: dimensions 1 and 2 in the pattern space were significantly correlated with STN axis1 and axis2 (arrows in the MDS plot), respectively (right). **(H)** Topographic organization of the IP_STN_. Left: GPe injection sites and IP_STN_ centroids superimposed onto coronal 3D views of the GPe and STN. Colors indicate the IP_STN_ centroid location along axis1, with the corresponding GPe inputs gradient direction (arrows). Right: correlations between GPe inputs and IP_STN_ along GPe gradient axes and STN axes. Correlations were calculated via Pearson’s r.

## Results

### Indirect pathway in the STN

To comprehensively analyze the organization of the indirect pathway in the mouse STN (IP_STN_), we employed a mesoscopic approach with the following components: 1) topographic labeling of subsets of GPe neurons with adeno-associated virus (AAV) expressing green or red fluorescent proteins (sfGFP or tdTomato), 2) computational reconstruction of axonal projection patterns, and 3) connecto-informatics of the GPe-STN circuit (**Figure 1; Figure S1–3;** see **STAR Methods**). To obtain IP_STN_ data, we conducted 73 injections into the GPe, serially imaged the GPe and STN, and then aligned our images to the Allen Mouse Brain Common Coordinate Framework (CCF v3, https://mouse.brain-map.org; **Figure 1A; Figure S1A; Movie S1**). After filtering our data to remove artifactual and weak projection signals, the results from 55 injections covering 84% of the GPe volume were further analyzed for IP_STN_ patterns (**Figure 1B; Figure S1B, C; Table S1**). Injections and axonal projections were reconstructed onto the segmented GPe and STN, respectively (**Figure S2**).

To investigate IP_STN_ connectivity patterns, we constructed heatmaps of GPe injection and STN projection (IP_STN_) distributions along the whole-brain anatomic axes, that is, anterior-posterior (AP), medial-lateral (ML), and dorsal-ventral (DV; **Figure 1C; Figure S3A, B**). The centrality of the signal distribution was used to identify injection and projection sites, and to subsequently measure their correlation. These heatmaps show that while projection signals were dispersed throughout the STN, injection signals were restricted to specific topographic loci in the GPe, which is desirable for fine connectivity mapping. Reconstructed STN projection patterns show a graded distribution with a notable degree of overlap, rather than divisible sub-territories (**Figure 1C–E; Figure S2, 3**) (Lambert et al., 2012; Mallet et al., 2007; Parent and Hazrati, 1995; Plantinga et al., 2018). This gradient was predominantly oriented along the ventromedial to dorsolateral axis, corresponding with the geometric longitudinal axis of the STN, termed “STN axis1”, and also along the transverse direction, perpendicular to the longitudinal axis, termed “STN axis2” (**Figure 1D, and 1F**). Although the centroid-based measurement effectively represents the projection sites, it may not fully portray the varying spatial pattern of broadly dispersed axonal projections. Therefore, we used multidimensional scaling (MDS) that visualizes the level of similarity of projection patterns in a 3D space, termed “pattern space” (see **STAR Methods**). Projections in a pattern space generated by MDS were continuously arranged with little or no noticeable segregation along two major dimensions which were significantly correlated with the STN axes (**Figure 1G**). Consistent with the centroid analysis, MDS confirmed graded topographic organization of IP_STN_ along the geometrical axes of the STN (STN axes 1 and 2). We then investigated the topographic relationships between inputs originating in the GPe and their projection targets by pairwise comparison of each GPe input locus and its corresponding projection locus in the STN along STN axes 1 and 2. Our analysis of input and projection distributions in the indirect pathway indicated that injection sites in the GPe are topographically aligned linearly along two directions, matching the geometric longitudinal and transverse axes of the GPe itself (**Figure 1H;** see **STAR Methods**). Interestingly, along STN axis2, the input-projection pattern was reversely related, revealing a “cross-over” projection pattern from the GPe to the STN. Our detailed analysis of GPe-STN connectivity clearly shows graded IP_STN_ organization along geometric axes, with generally extensive overlaps and an absence of sharp boundaries.

### Hyperdirect pathway in the STN

Next, to comprehensively assess the organization of the hyperdirect pathway in the STN (HP_STN_), we extracted data from the Allen Mouse Brain Connectivity Atlas dataset (AMBCA, https://connectivity.brain-map.org) (Oh et al., 2014) regarding axonal projections following isocortical injections. Our data-filtering criteria included detectable STN projection signals and contamination of cortical injection (**Figure 2A; Figure S1A and 4A–C;** see **STAR Methods**). The resulting dataset included 176 injections with AAV-expressing fluorescent proteins throughout the isocortex, covering 71% of the volume of layer (L) 5 in wild-type C57BL/6J mice and nine transgenic Cre lines shown to have L5-specific expression of Cre (**Figure 2B; Table S2, 3**). Additionally, our own retrograde tracer experiments using retro-AAV or -Lenti systems (Kato et al., 2014; Tervo et al., 2016) confirmed that HP_STN_ arises in various cortical subregions, mainly from a subpopulation of corticofugal neurons in the ipsilateral L5, which is consistent with previous observations (Kita and Kita, 2012) (**Figure S4D**).

**Figure 2.**
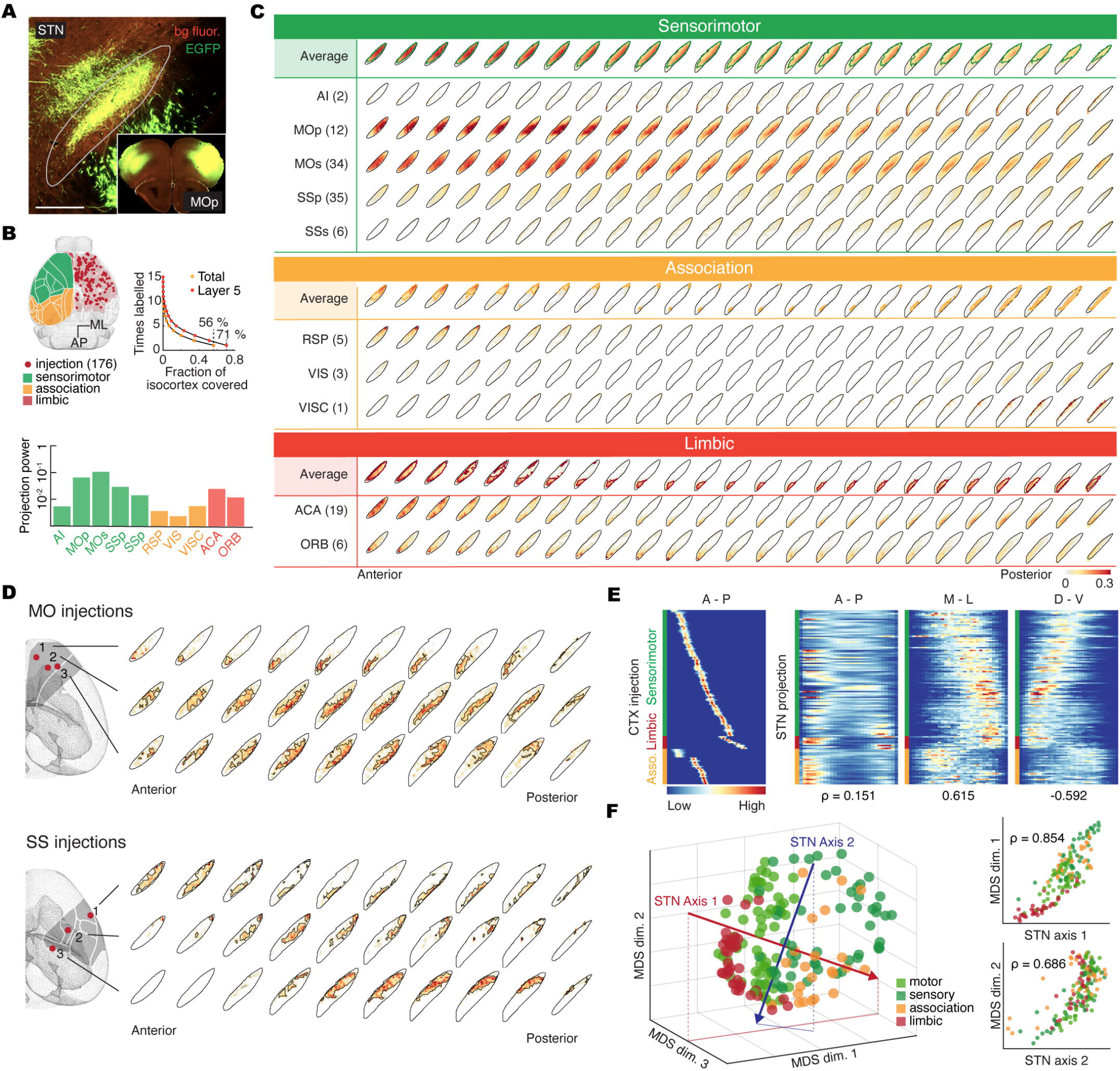
Graded topography of hyperdirect pathway in STN. **(A)** Representative axonal projections in the STN (HP_STN_) from an MOp injection (inset) extracted from AMBCA datasets. Scale bar, 250 µm. **(B)** Top: coverage (left) and spatial locations of the hyperdirect cortical injection site (right) across the isocortex (n = 176). Bottom: Bar plot of STN projection power for each subregion, an estimated STN projection volume assuming the subregion is fully labelled, shown in log-scale. Grouped functional parts of cortical subregions are color-coded (See **Table S3** for functional classification). **(C)** Digitally reconstructed serial STN projection density patterns of the averaged HP_STN_ from each grouped functional part and constituting cortical subregion at 25-µm intervals. The number in parentheses denotes the number of localized injections within the subregion. Subregions without localized injections are not shown. **(D)** Representative set of intra-subregion injections within the MO (top) and SS (bottom), and corresponding serial STN projection density at 50-µm intervals. **(E)** Heatmap showing cortical inputs and HP_STN_ signal distribution along anatomical axes (columns). Cortical injection experiments (rows) were grouped into three functional parts and sorted according to the AP of the injection centroid in each group. **(F)** Left: HP_STN_ projection patterns in the pattern space (same as in Figure 1G). Right: Dimensions 1 and 2 in the HP_STN_ pattern space were also significantly correlated with STN axis1 and axis2 (arrows in the MDS plot). Correlations were calculated by Pearson’s r.

Information flow from the cortex to the BG is believed to progress through a hierarchical series of sensorimotor, association, and limbic areas, supporting the tripartite STN model (Lambert et al., 2012; Mallet et al., 2007; Parent and Hazrati, 1995; Plantinga et al., 2018). However, much of the data leading to this view regarding the subdivisions of the STN are qualitative, variable, fragmented, and difficult to reconcile, owing to the technical limitations of neuroanatomical research (Alkemade and Forstmann, 2014; Keuken et al., 2012; Lambert et al., 2015). Therefore, to update this view of the STN, we first performed an inclusive analysis at the level of region-to-region connectivity of the HP_STN_. We found that STN projection patterns from each cortical subregion exhibited distinct yet extensively overlapping projection patterns (**Figure 2C**). For instance, projections from motor cortical areas such as the primary and secondary motor areas (MOp and MOs) broadly targeted the center region of the STN, while those from sensory areas such as primary and supplemental somatosensory areas (SSp and SSs) targeted the dorsolateral STN (anatomic abbreviations and functional groups shown in **Table S2**). Association areas, such as the retrosplenial (RSP), visual (VIS), and visceral (VISC) area, and limbic areas, such as anterior cingulate (ACA) and orbital (ORB) area, contained dorsolateral and ventromedial STN projections spanning the central part of STN, respectively. Overall, our analysis of STN projection patterns showed much broader coverage, with more extensive overlap than previously thought (Lambert et al., 2012; Mallet et al., 2007; Parent and Hazrati, 1995; Plantinga et al., 2018). Furthermore, within motor and sensory cortices, we found that individual injections revealed variable and distinct yet overlapping projections in the STN (**Figure 2D**). Hence, we investigated the organization of the HP_STN_ using non-biased techniques, such as a voxel-based analysis that disregards anatomically defined cortical subregions. Similar to our analysis of the IP_STN_ described above, we examined the distributions of cortical inputs and STN projections along the whole-brain anatomic axes (AP, ML, and DV). We found that the HP_STN_ had a graded distribution with a notable degree of overlap along STN axis1 (**Figure 2E**). Furthermore, MDS analysis indicated that the 3D projection patterns of the HP_STN_ were gradually distributed along STN axis1 and axis2 (**Figure 2F**). Together, these results indicate that the HP_STN_ has graded organization along geometric axes, with overlaps and an absence of sharp boundaries, which prompts re-evaluation of the conventional view of a functionally subdivided STN determined by cortical input patterns.

### Hyperdirect pathway in the GPe

We expanded our analysis of the hyperdirect pathway to include projection patterns in the GPe. Based on previous reports (Abecassis et al., 2020; Karube et al., 2019; Naito and Kita, 1994; Winnubst et al., 2019) showing the existence of direct cortico-pallidal connections, we questioned whether the hyperdirect pathway providing direct cortico-pallidal projections into the GPe (called HP_GPe_) collaterally provides HP_STN_, and how HP_GPe_ displays topographic distributions. First, we injected retro-AAV-iCre and Cre-dependent AAV-JxON-tdTomato into the GPe and MOp, respectively, and then cleared the tissue. Our results revealed strong collateral projections from labeled MOp neurons in both the GPe and STN with additional downstream targets in the substantia nigra (SN) and superior colliculus (SC), but few or no targets in contralateral cortical areas (**Figure 3A**). As with the HP_STN_, we observed graded organization within the HP_GPe_. The HP_GPe_ appeared to be topographically organized with fidelity to the whole-brain anatomic axes and segmented into a tripartite form. This was especially the case with respect to inputs arising in association and limbic areas, which is comparable to the HP_STN_ (**Figure 3B**). This organization might arise through the innate geometric situation of the GPe matched with the whole-brain anatomic axes and/or the larger receptive volume of the GPe compared with the STN. Next, we conducted pairwise topographic matching between the loci of cortical inputs and projections in the GPe (HP_GPe_). We found two major axes of topographic organization of hyperdirect pathway inputs and the HP_GPe_ that were correlated and reversely correlated along STN axis1 and axis2, respectively (**Figure 3C, D; Figure S5A**). These gradient axes of the HP_GPe_ appeared to match the geometric axes of the GPe, which is similar to the gradient axes observed for input into the indirect pathway. To examine the relationship between the two pathways along these gradient axes, we performed a paired comparison between the IP and HP patterns. For this analysis, we measured the similarity between the pathways in the STN and GPe, respectively. The HP_STN_ and IP_STN_ projection patterns were used to examine the gradient similarity in the STN, while the HP_GPe_ and indirect pathway input (IP_INPUT_) were used for that in the GPe. We then calculated the correlation between these pairwise similarities in the STN and GPe. We found strong similarities in STN projection patterns between the IP_STN_ and HP_STN_ when the IP_INPUT_ and HP_GPe_ locus were located close together (**Figure 3E**).

**Figure 3.**
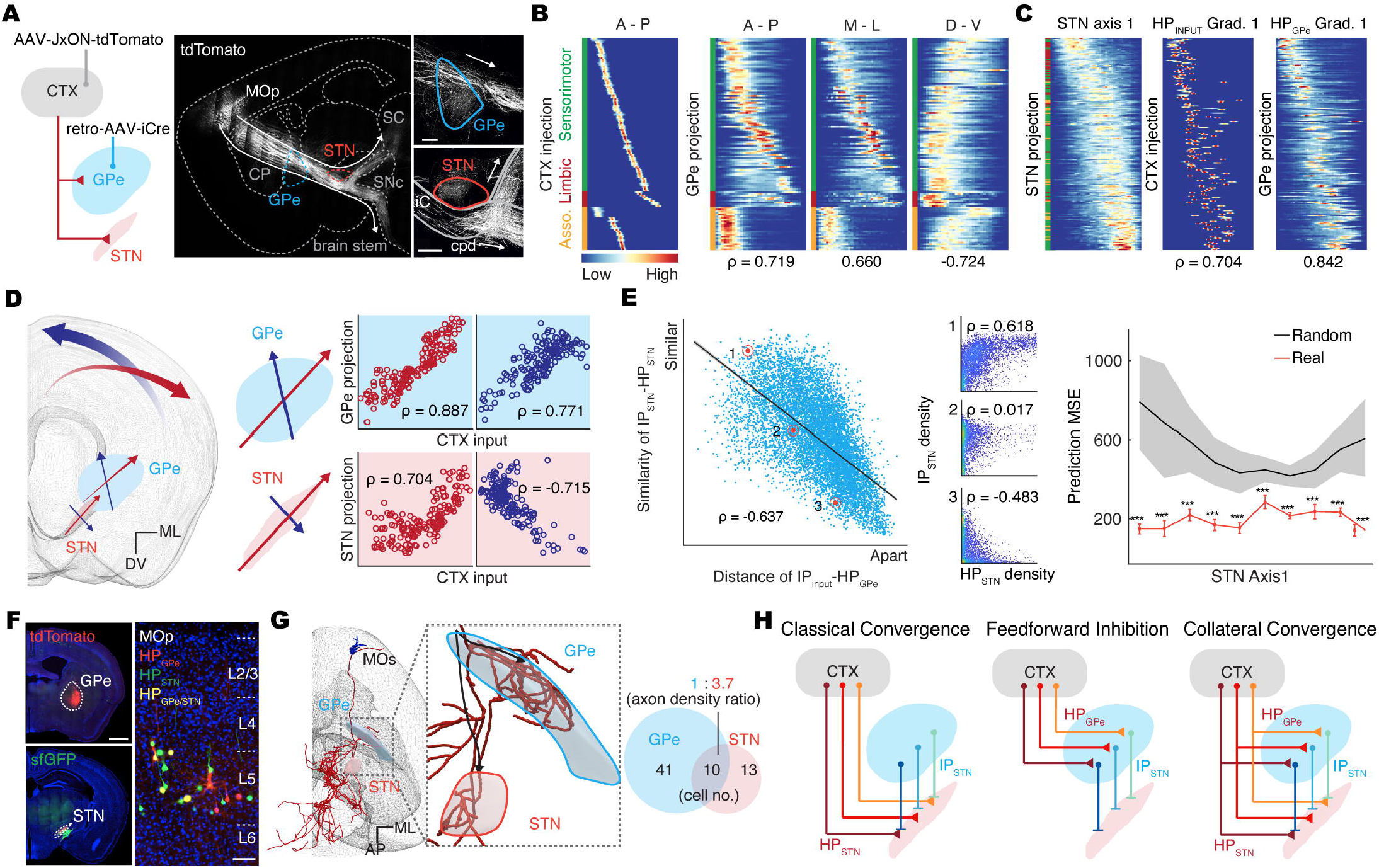
Complex integration of indirect and hyperdirect pathways in STN. **(A)** Left: illustration of the hyperdirect pathway with a direct pallidal connection and schematic of the injection strategy for labelling collateral hyperdirect projections (retroAAV-iCre and Cre-dependent switch-on AAV-tdTomato were injected in the GPe and MOp, respectively). Right: z-projection image of a cleared parasagittal section (2.5 mm thick) showing hyperdirect projections both in the GPe and the STN, and the trajectories of the HP with two downstream targets (arrows). Scale bar, 250 µm. **(B)** Heatmap showing cortical inputs and HP_GPe_ signal distribution along anatomical axes (columns). Cortical injection experiments (rows) were grouped into three functional parts and sorted according to the AP of the injection centroid in each group. **(C)** Heatmap of sorted HP_STN_ with cortical inputs and HP_GPe_ signal distribution along the STN axis1 and corresponding cortex and GPe gradient axis. Rows are sorted by HP_STN_ axial location along STN axis1. **(D)** Left: illustration of the topographic organization of the HP_STN_, HP_GPe_, and cortical inputs with STN geometric axes and corresponding gradient axes of the HP_GPe_ and cortical inputs (arrows). Right: linear relationship between axial locations of the HP_GPe_ (top), HP_STN_ (bottom), and cortical inputs. **(E)** Topographic relationship between the IP and HP. Left: the projection similarity between IP_STN_-HP_STN_ was inversely correlated with the distance between IP_INPUT_-HP_GPe_. Each dot represents a pair of IP and HP experiments (n = 55 × 176). Middle: Scatterplots of IP_STN_ and HP_STN_ density from three representative pairs showing similar (1), intermediate (2), and different (3) IP_STN_ and HP_STN_ projections. Right: Prediction of HP_GPe_-HP_STN_ based on GPe input-IP_STN_ topographic organization resulted in significantly lower mean squared error (MSE) compared with randomizations (***p < 0.001, two-sided Student t-test). Red error bars represent the mean with s.e.m, while the gray area represents the 5^th^ to 95^th^ percentile of randomization. **(F)** Left: Injection sites of retroAAV expressing tdTomato and sfGFP in the GPe (top) and STN (bottom), respectively. Right: Magnified image of the MOp showing labeled GPe (red)- and STN (green)-projecting cells in L5, with co-labeled (yellow) cells representing collateral GPe/STN-projecting neurons. Scale bar 1 mm and 100 µm (enlarged view). **(G)** Analysis of single cell reconstruction data from the MouseLight database. Left: representative MOs neuron (AA0772) collaterally projecting to the GPe and STN. Right: cortical neuronal population innervating the HP_GPe_ (64%), HP_STN_ (20%), and collaterally innervating the HP_GPe/STN_ (15%). **(H)** Schematic illustration of three types of signal integration in the STN from indirect and hyperdirect pathways. Correlations were calculated via Pearson’s r.

### Complex integration of the indirect and hyperdirect pathways in the STN

Given our findings that the IP_STN_, HP_STN_, and HP_GPe_ are topographically organized with gradient convergence at the mesoscopic level, we next performed a detailed analysis of HP connectivity at the cellular level. To dissect HP connectivity types, we first labeled cortical neurons innervating HP_STN_ and/or HP_GPe_ by injecting retro-AAV-tdTomato and -sfGFP in the STN and GPe, respectively. We identified three types of cortical neurons in L5 of MOp: Those innervating only HP_STN_, only HP_GPe_, and both HP_GPe/STN_ (**Figure 3F**). Additionally, by data-mining single-neuron reconstruction data from the MouseLight project (http://www.mouselight.janelia.org) (Winnubst et al., 2019), we confirmed the existence of these three types of cortical neurons (**Figure 3G; Figure S5B, C**). Neurons innervating the HP_STN_ are predominantly pyramidal tract (PT) neurons, while those innervating the HP_GPe_ are both PT and intratelencephalic (IT) neurons, with a subset (∼15 %) that collaterally innervate the HP_GPe/STN_. Notably, the axonal density of neurons innervating the HP_GPe/STN_ in the STN is 3.7-fold higher than that in the GPe. Consistent with our mesoscopic retrograde tracer data, shown in **Figure 3A**, we found that cortical neurons innervating the HP_GPe/STN_ send collateral innervations to multiple brain areas such as the striatum (STR), SN, SC, and others, which suggests that these neurons may help orchestrate movement. Although we analyzed a small set of reconstructed single neurons, their projection patterns were consistent with our mesoscopic, topographic analysis of the HP_GPe_ and HP_STN_ (**Figure S5D**). Our results indicate that at least three types of signals from the indirect and hyperdirect pathways in the STN are integrated, including a classical form of convergence of inhibition and excitation, feedforward inhibition, and collateral convergence (parallel after-discharge-like circuitry; **Figure 3H**). In addition, we confirmed HP projections into the human STN at the macroscopic scale by 7T-MRI-based *in vivo* ultrahigh-resolution fiber tractography of a healthy human brain (**Figure S6; Table S4**; see **STAR Methods**). Our seed-based probabilistic tractography of the HP revealed a direct cortico-pallidal connection, the HP_GPe_, and a collateral HP_GPe/STN_ in the human brain, similar to those we observed in mice (**Figure S6B, C**). We divided cortical subregions into three groups: motor, limbic, and association (further divided into prefrontal- and rest-association) areas (**Table S2**). HP projections from the association, motor, and limbic areas were only somewhat segregated in the ventromedial, dorsolateral, and ventrolateral areas, respectively, appearing along the longitudinal axis of the STN (axis1). However, as we observed in the mouse HP_STN_, there was a high degree of overlap and no evident anatomical borders or clear subdivisions (**Figure S7D–H**). We also found the HP_STN_, HP_GPe_, and collateral HP_GPe/STN_ organization in the human brain to be consistent with the complex signal integration patterns in the mouse. In agreement with our mouse connectivity maps, our analysis of human 7T MRI-based tractography data revealed considerable and gradual overlap along the longitudinal axis of the human STN. Together, these results suggest that graded convergence of IP_STN_ and HP_STN_ is organized along the STN geometric axes, and that GPe neurons innervating the IP_STN_ might be activated by the same HP_STN_. These circuit features imply functional computational complexity and delicate control, updating classical models (DeLong and Wichmann, 2015) with the idea that the indirect and hyperdirect pathways participate in relaying and integrating information needed for coordinated movement.

### Cellular composition of the STN

The way in which STN neurons process convergent inputs from the IP and HP depends critically on the molecular and cellular properties of these neurons. The STN is generally described as a homogenous glutamatergic nucleus of the BG. To test this, we used serial single-molecule fluorescence in situ hybridization (smFISH) with a custom computational analysis for cell-type markers and receptors to examine the topographic molecular and cellular profile of the STN (**Figure 4A**; see **STAR Methods**). Most STN neurons identified by the neuronal marker *Rbfox3* co-expressed the glutamatergic marker *Slc17a6*, while very few neurons co-expressed the GABAergic marker *Slc32a1*. Approximately half of the STN cells expressed glia-associated genes (*Aqp4*, *Mbp*, and *Tmem119*) (**Figure S7**). Thus, our serial smFISH, immunostaining, and topographic analysis confirmed that most neurons in the STN are purely glutamatergic. Previous anatomical studies have reported the presence of parvalbumin-expressing (PV+) neurons in the STN (Alkemade et al., 2019; Hontanilla et al., 1998; Lévesque and Parent, 2005; Wu et al., 2018). PV expression is conventionally interpreted to reflect a GABAergic subpopulation (Hu et al., 2014), which is inconsistent with our serial smFISH results and previous studies (Lévesque and Parent, 2005; Roccaro-Waldmeyer et al., 2018). Accordingly, we conducted detailed cellular characterization of PV+ STN neurons. First, we performed triple smFISH for *Pvalb*, *Slc17a6*, and *Slc32a1*. We found that *Pvalb*+ neurons in the STN exclusively co-express glutamatergic marker *Slc17a6*, but not GABAergic marker *Slc32a1*, whereas those in the other brain areas such as hippocampus, cortex, and zona incerta co-express *Slc32a1* (**Figure 4B–D; Figure S8A**). Notably, quantitative analysis of the topographic distribution of *Pvalb*+ neurons indicated that glutamatergic PV+ neurons were particularly prevalent in the dorsolateral and middle STN (**Figure 4C, D**). Our serial multiplex-smFISH data provide direct evidence that PV+ STN cells are glutamatergic with topographic segregation in the STN (Hontanilla et al., 1998; Wu et al., 2018) and suggest that parvalbumin cannot reliably serve as a marker for GABAergic interneurons, together with previous findings (Jinno and Kosaka, 2004; Shang et al., 2015; Wallace et al., 2017).

**Figure 4.**
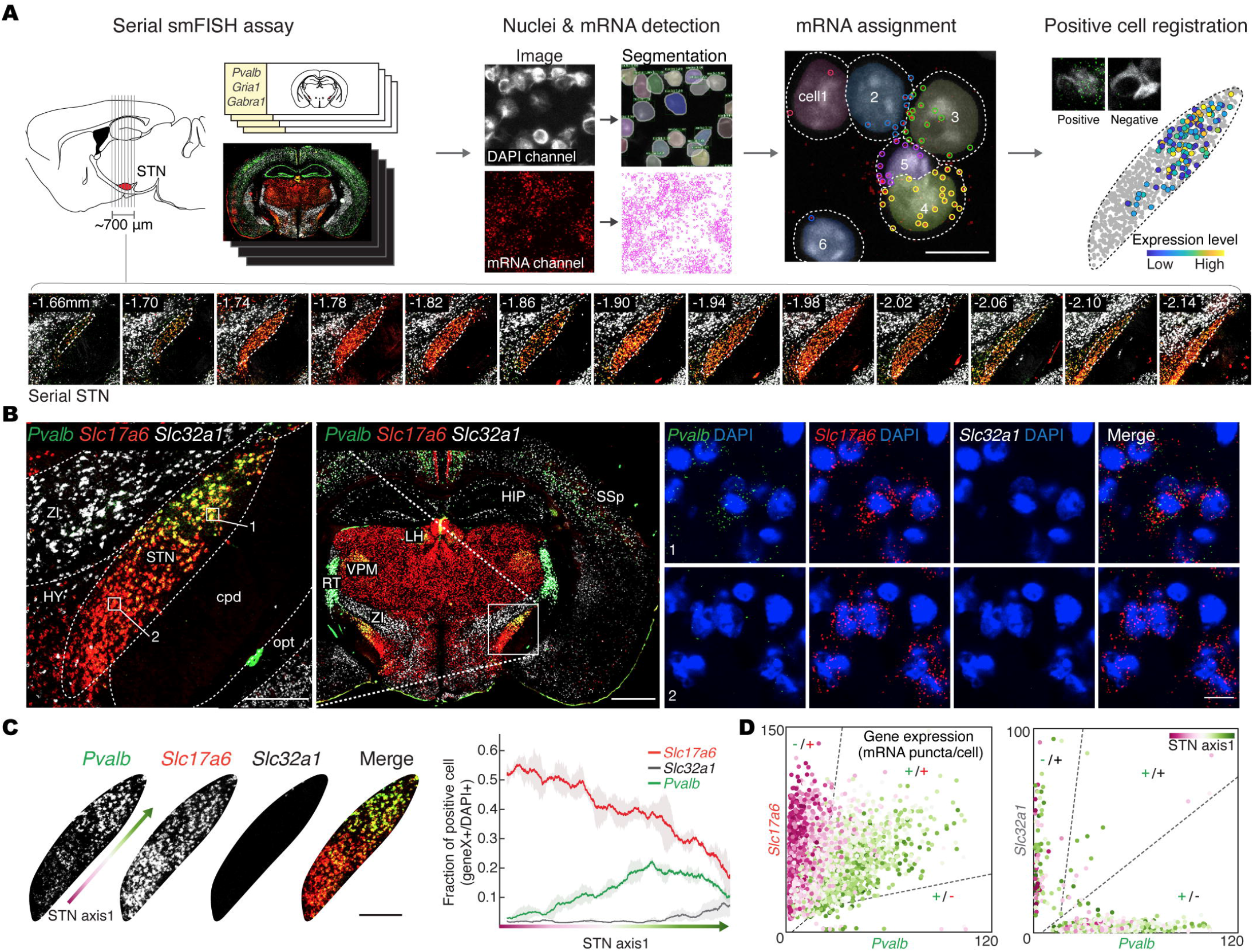
Glutamatergic STN PV+ neurons. **(A)** Top: pipeline of serial smFISH data generation and processing. Bottom: example serial smFISH image of STN with the distance from the bregma indicated at the corner. **(B)** Left: example smFISH image of the STN (inset) from coronal brain slice with *Pvalb*, *Slc17a6*, and *Slc32a1*. Right: magnified views of the regions indicated by white boxes in the dorsolateral (top) and ventromedial (bottom) STN (left panel) show that STN neuronal population co-express *Pvalb* and *Slc17a6*, but not *Slc32a1*. Scale bar, 1 mm (left), 250 µm (left, inset), and 10 µm. **(C), (D)** Topographic distribution of glutamatergic PV+ neurons in the STN. Representative smFISH images cropped to show the STN (**C**, left), and quantification of cells expressing each gene along STN axis1 shown with the mean (n = 2) and s.e.m. (**C**, right). The pseudocolor arrow represents STN axis1. Co-expression plot of *Pvalb* with *Slc17a6* (**D**, left) and *Slc32a1* (**D**, right), showing the number of mRNA puncta per cell for each gene. Each dot represents a cell, colored according to location along STN axis1. Dashed lines represent the thresholds for positive cells. Double negative cells are not shown. Scale bar, 250 µm.

To examine further physiological properties of PV+ STN neurons, we used PV-IRES-Cre::Ai6 mice expressing ZsGreen in PV+ cells. We first confirmed that PV+ neurons in the STN of these mice were immunohistochemically positive for glutamate and negative for GABA (**Figure 5A**). By clearing parasagittal sections (2.5 mm in thickness) that embrace the entire STN without making physical cuts, we 3D-mapped total PV+ neurons in the STN and found topographically segregated distribution in the dorsolateral and middle STN, consistent with serial-smFISH data (**Figure 5B; Movie S2**). Because Parkinsonian symptoms are correlated with an increase in burst firing in STN neurons, we next recorded the firing patterns of glutamatergic PV+/- STN neurons using PV-IRES-Cre::Ai6 mice. Both PV+ and PV− neurons displayed similar spontaneous firing and other basic properties including input resistance (**Table S5**). However, only PV+ neurons showed phasic burst firing, which we refer to a firing pattern, composing of fast firing (an interspike interval, ISI, of less than 10 ms) and a pause; even current injection could not evoke bursts from PV− neurons (**Figure 5C-F; Figure S8B-E; Table S6;** see **STAR Methods**). Burst and tonic firing patterns were observed at an equivalent membrane potential range (−58 to −45 mV) (**Figure S8E**), suggesting that burst firing was not due to differences in the membrane potential *per se*, even though firing patterns in STN neurons can be affected by membrane potentials (Kass and Mintz, 2006). Injection of negative current pulses evoked significantly smaller hyperpolarization-induced depolarizing sags in PV+ vs. PV− neurons, which was consistent with their burst properties (**Figure 5F**). Taken together, our results provide clear evidence that there are at least two functional types of glutamatergic neurons in the STN, and that glutamatergic PV+ neurons contribute to burst firing in the dorsolateral and middle STN. This is compatible with previous studies of spatial distribution of firing patterns by intraoperative microelectrode recording within the human STN showing the occurrence of burst firing in in the dorsal STN while tonic firings towards the ventral STN (Kaku et al., 2019).

**Figure 5.**
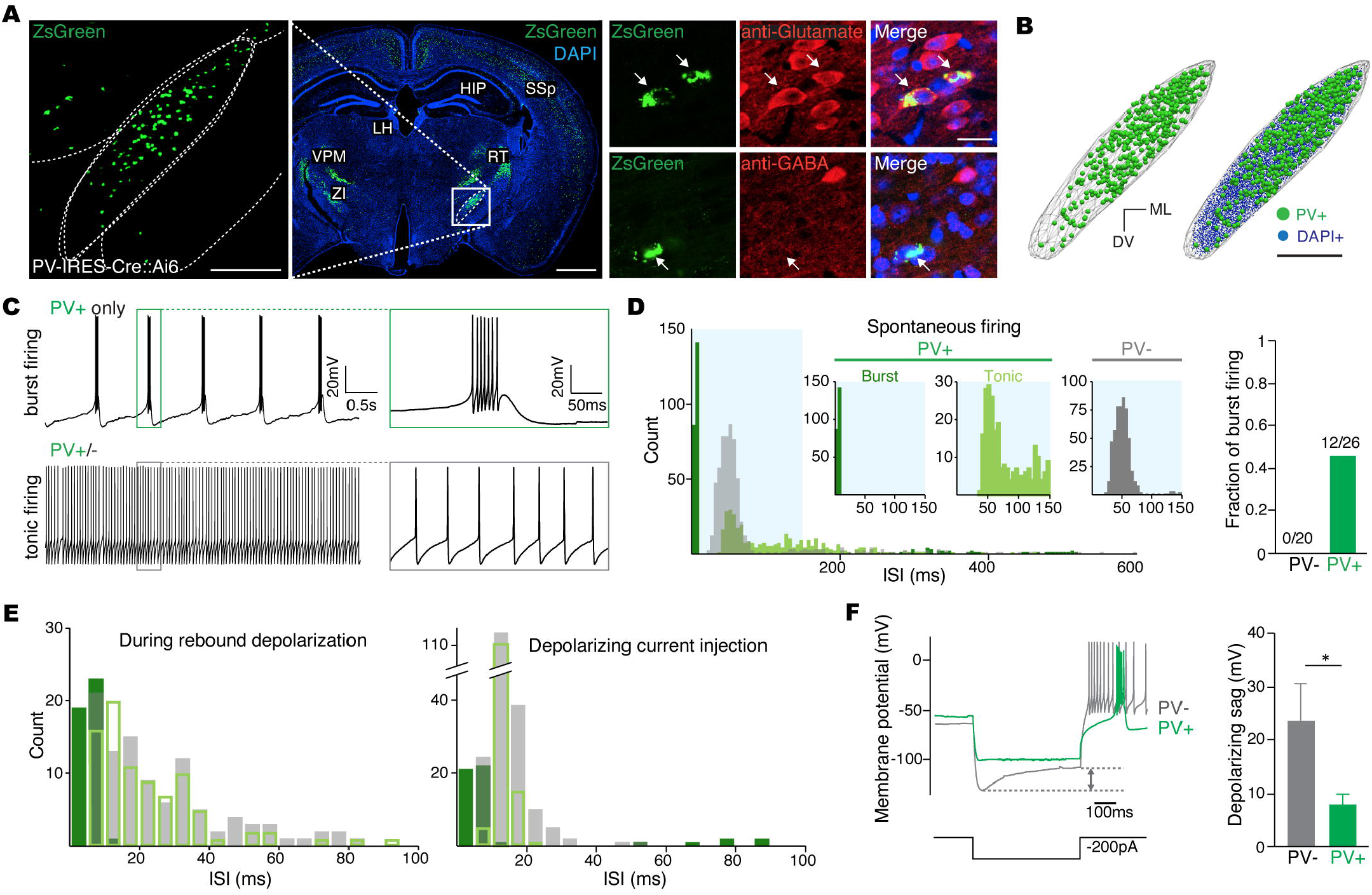
Firing patterns of STN PV+/- neuron. **(A)** Left: representative image of the STN (inset) from coronal brain slice of PV-IRES-Cre::Ai6 mouse line, showing PV+ neurons. Right: magnified immunofluorescence image of STN cells show STN PV+ neurons co-labeled with anti-Glutamate (top) but not with anti-GABA (bottom). Scale bar, 1 mm (left), and 250 µm (left, inset) and 20 µm (right). **(B)** Distribution of PV+ neurons in the STN using automatic cell detection from PV-IRES-Cre::Ai6 mouse. Scale bar, 250 µm. **(C)** Electrophysiological properties of STN PV+/-neurons. Representative tonic (bottom) and burst (top) firing with enlarged traces shown in boxes, recorded from STN PV+/- neurons using the PV-IRES-Cre::Ai6 mouse line. Interspike interval (ISI) of first two, 2^nd^ and 3^rd^, last two spikes, and average of all spikes shown in the boxes; 7.7, 5.9, 7.7, and 6.5 ms for PV+ burst firing; 50.3, 51.4, 46.7, and 47.5 ms for PV-tonic firing neuron. **(D)** ISI histogram of spontaneous firing patterns (burst PV+ and tonic PV+/- neurons, n = 12, 14, and 20, respectively). Insets in the top panels show magnified ISI histogram during 0-150 ms (shaded area). Bar graph shows the proportion of burst firing cells (right). **(E)** ISI histogram of firing patterns during a rebound depolarization following the removal of a hyperpolarizing current injection (left), and during depolarizing current injection (right; see **Methods**) **(F)** The hyperpolarization-induced depolarizing sag of PV+/- neurons, marked by an arrow (left), was significantly smaller than that in PV+ neurons (right). Data are mean ±s.e.m. Mann–Whitney U test with p < 0.01. ISI of first two, 2^nd^ and 3^rd^, last two spikes, and average of all spikes; 4.2, 3.5, 8.9, and 4.6 ms for PV+ burst firing, 12.4, 12.5, 55.1, 24.2 ms for PV-tonic firing neuron.

### Cell type-dependent synaptic connectivity

Our analysis defined new functional cell types and established wiregrams for the IP_STN_ and HP_STN_, allowing us to refine our analysis of synaptic connectivity in the IP and HP to be cell type-specific. Thus, we tested whether the IP_STN_ and HP_STN_ form PV+/- cell-specific synaptic connections, and whether the HP_GPe_ forms synapses on the two major GABAergic cell types in the GPe, that is, prototypic neurons projecting to the STN and arkypalidal neurons projecting to the striatum (Mallet et al., 2012; Mastro et al., 2014). By combining retrograde virus injections and mammalian GFP reconstitution across synapses (mGRASP) (Kato et al., 2014; Kim et al., 2012), which is an advanced method for accurately detecting synaptic connections, we mapped synaptic connectivity of these four cell-types in IP_STN_, HP_STN_, and HP_GPe_ (**Figure 6A-F**; see **STAR Methods**). We identified synaptic connections in STN PV+ and PV− neurons originating in both the IP_STN_ and HP_STN_, by injecting AAV expressing the pre- and post-synaptic mGRASP components into the GPe (for IP_STN_), cortex (for HP_STN_), and STN, respectively. STN PV+/- neurons were distinguished by injection of AAV expressing iRFP in a Cre-dependent manner into the PV-Cre mouse line. We also observed synaptic connections in both GPe prototypic and arkypalidal neurons (labeled by a combination of retro-Lentivirus expressing iCre injected into the STN and STR, respectively, and AAV expressing Cre-On-postsynaptic mGRASP into the GPe) projecting from the HP_GPe_ (labeled by injecting presynaptic mGRASP in the cortex). Our analysis revealed that axonal projections in both the STN and GPe from the IP_STN_ and HP_STN_, and from the HP_GPe_, respectively, indeed serve as inputs. Furthermore, our results indicated that all four cell types in the STN and GPe (STN PV+, PV-, GPe arkypalidal, and prototypic neurons) communicate with each other and receive synaptic information from the cortex. Thus, in terms of cortex-GPe-STN circuitry, our data indicate that “everyone talks and everyone listens.” Next, in terms of connectivity patterns, the two types of STN neurons integrate excitatory and inhibitory signals (E/I) from the IP_STN_ and HP_STN_. We examined the differential expression of glutamate and GABA receptors in STN PV+/- neurons. To this end, we performed serial triple smFISH for *Gria1* (encoding GluA1), *Gabra1* (GABA_A_ α1 subunit), and *Pvalb*, and quantitatively compared their expression with respect to the topographically graded innervations of the IP_STN_ and HP_STN_ (**Figure 6G-I**). Our co-expression plot of *Gria1* and *Gabra1* showed a strong correlation between these genes, indicating that most STN neurons integrate both excitatory and inhibitory inputs at the single-cell level. Notably, the total expression levels of *Gria1* and *Gabra1* formed a gradient along the STN axis1 with higher and lower expression in the ventromedial and dorsolateral STN, respectively, similar to the expression pattern of *Slc17a6* (**Figure 4C, and 6H, I**). At the cellular level, the *Gria1* to *Gabra1* expression ratio was approximately similar in the dorsolateral PV+, but this expression ratio was significantly higher in ventromedial PV− neurons (**Figure 6I**). Taken together, our results indicate that two types of glutamatergic PV+/- STN neurons, which integrate information from both the IP_STN_ and HP_STN_, and fire in different patterns, are segregated along a topographical distribution and express different ratios of genes encoding E/I receptors. Although further direct evidence is needed, these findings may represent a step forward in characterizing the E/I imbalance of BG circuits in Parkinson’s disease models, as well as the effects of DBS. Interestingly, when we analyzed DBS results with respect to electrode position within specific STN structures from 26 Parkinson’s disease patients who had received bilateral STN DBS implementation together with previous studies (Herzog et al., 2004; Mallet et al., 2012; Wodarg et al., 2012), we found that patients showing DBS-associated movement improvement had electrodes placed in the dorsolateral and middle STN, with somewhat greater improvement associated with electrode placement towards the middle of STN axis1 (**Figure S9**). This part of the STN has been reported to receive direct inputs from motor cortical areas and to contain PV+ glutamatergic neurons (Alkemade et al., 2019; Eisenstein et al., 2014; Herzog et al., 2004; Lévesque and Parent, 2005; Mallet et al., 2007; Wodarg et al., 2012). The correlation among position-dependent STN DBS effects and connectivity patterns led us to hypothesize that distinct features of connectivity and cellular composition in the dorsolateral and middle STN influence the impact of DBS.

**Figure 6.**
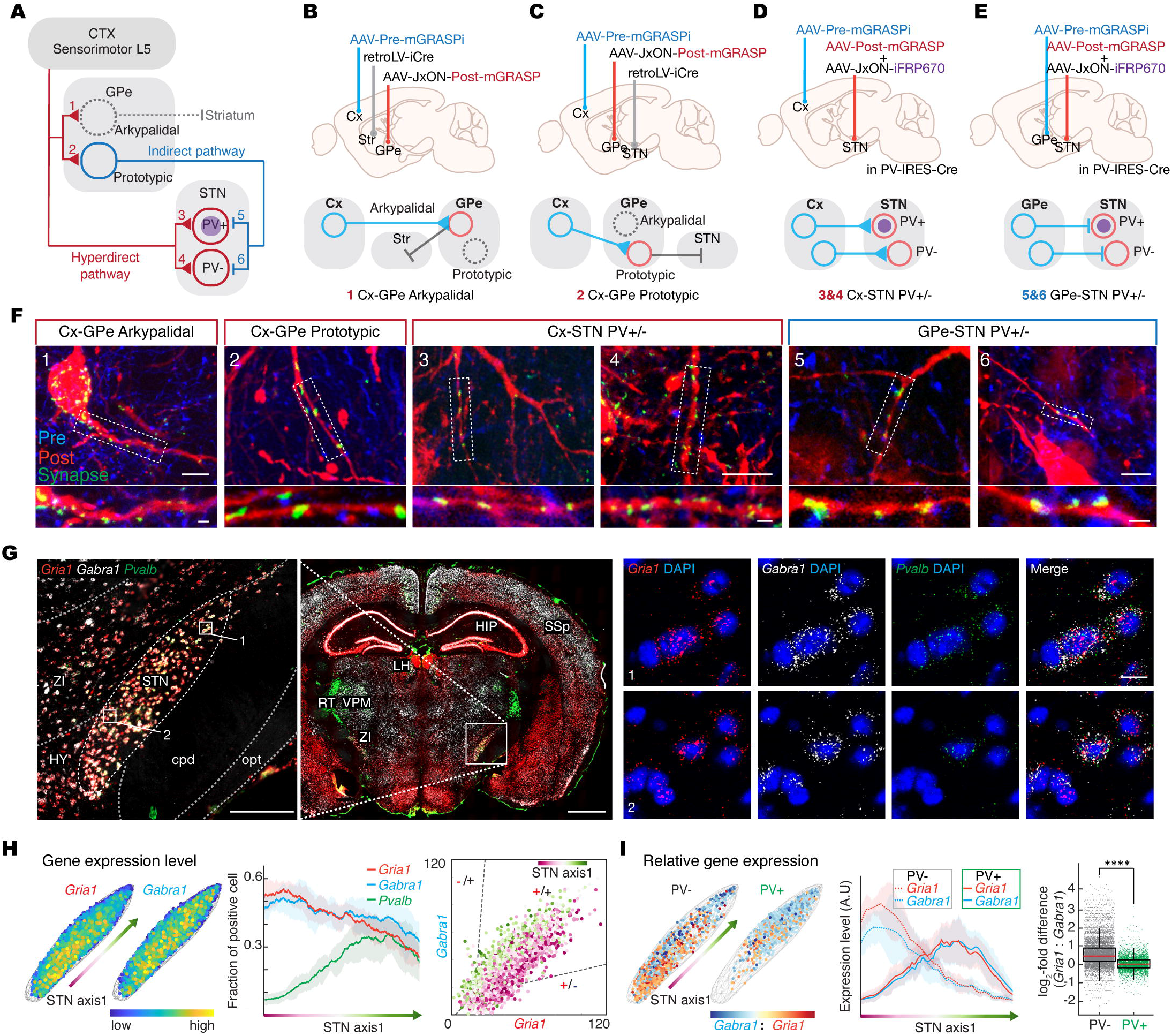
Cell type-dependent synaptic connectivity. **(A)** Scheme of tested cell type-specific connections in the HP_GPe_ (1, 2), HP_STN_ (3, 4), and IP_STN_ (5, 6). **(B-E)**, Labeling strategies for testing synaptic connections listed in **(A)** using retrograde virus and mGRASP system. Presynaptic-mGRASP was injected into either the cortex (1–4) or the GPe (5–6). Projection-dependent GPe cell types (arkypallidal and prototypic) were labeled with retro-lenti virus expressing Cre in the Striatum and STN, respectively (1, 2), followed by Cre-ON postsynaptic-mGRASP in the GPe. A mixture of postsynaptic-mGRASP and Cre-ON fluorescent protein was injected into the STN of PV-IRES-Cre mice to distinguish STN PV+/- neurons (3–6). **(F)** Post-synaptic neurons (top) and enlarged image of dendrite (bottom), marked by white box. Green mGRASP signals along the dendrites of post-synaptic neurons (red) were detected in all tested sets, indicating the synaptic connections. Scale bar, 10 µm (top), and 2 µm (bottom). **(G)** Representative coronal smFISH image of the STN with *Pvalb*, *Gria1,* and *Gabra1*. Enlarged views of the regions indicated by white boxes in the dorsolateral (top) and ventromedial (bottom) STN (left panel) showing co-expression of *Gria1* and *Gabra1*. Scale bar, 1 mm (left), 250 µm (left, inset) and 20 µm (right). **(H)** Topographic distribution along STN axis1 of cells expressing *Gria1* and *Gabra1.* Quantification of cells expressing each gene along STN axis1 (left) and a co-expression plot (right) showing that most STN cells co-express *Gria1* and *Gabra1*. Data are represented as means (n = 6) with s.e.m. **(I)** Left: Relative expression level of *Gria1* and *Gabra1* in PV+/- neurons. Middle: *Gria1* to *Gabra1* ratio in PV+/- neurons exhibit distinct topographic pattern along STN axis1. Right: the *Gria1* to *Gabra1* ratio was significantly lower in STN PV+ cells (permutation test, n = 10,000, ****p < 0.0001). Data are represented as means (n = 6) with s.e.m.

In Sum, our comprehensive profiling of circuit- and cellular-level connectivity of the indirect and hyperdirect pathways revealed a topographically graded organization with three convergent types of indirect and hyperdirect-pathway (GPe-only, STN-only, and both) in the STN. Topographically distinct molecular, cellular, and physiological properties together with the circuit features imply functional computational complexity and delicate control. Our results suggest a complex interplay of information within the basal ganglia further on than the general consensus of parallel segregated input patterns with homogeneous cytoarchitecture and physiology. Our new view of the indirect and hyperdirect pathways, cytoarchitecture, and physiology properties substantially updates classical models and suggests, from a structural basis, a way to view relayed and integrated information flow underlying coordinated movement controls and STN DBS effects (**Figure 7**).

**Figure 7.**
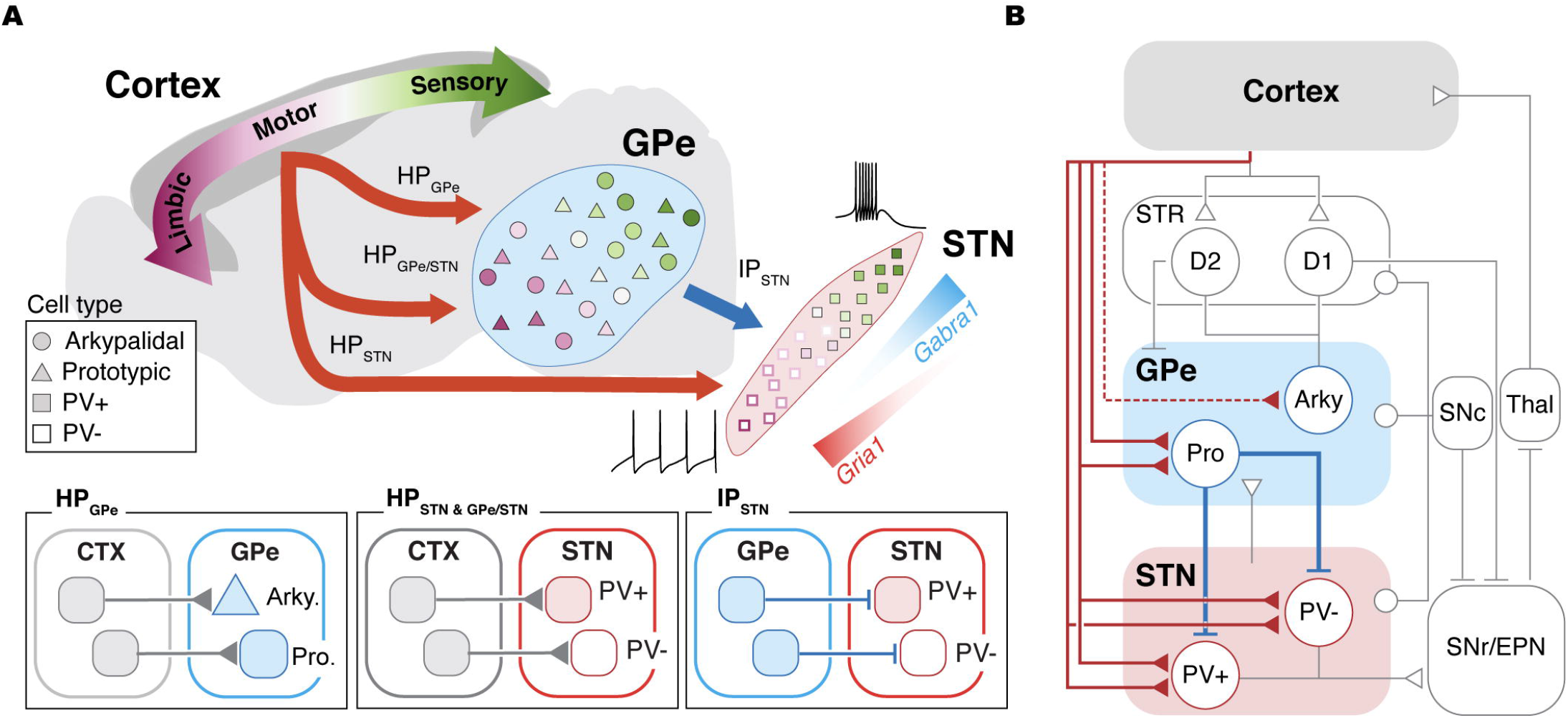
Schematic of basal ganglia network with detailed topographic connectivity, cytoarchitecture, and firing pattern of the STN. **(A)** Summary of connectivity, cellular and physiological profiles (top) and cell-type specific synaptic connectivity (bottom) of the STN. Topographical organization of three convergent types of indirect and hyperdirect pathways is represented in gradual color change and arrows. Representative traces of two firing patterns, burst and tonic firing, illustrate topographic distribution of functionally distinct cell-types along dorsolateral and ventromedial parts of the STN, respectively. **(B)** Schematic of basal ganglia network with updated circuit- and cellular-level connectivity found in this study (represented in colored line). Three convergent hyperdirect and indirect pathways related to the STN are shown in solid lines.

## Discussion

Understanding how the STN controls motor functions, and identifying the circuit-level mechanisms underlying the effects of STN-DBS requires detailed convergent connectivity profiles for the main inhibitory and excitatory inputs in the STN. Furthermore, molecular profiles of cell types and neurotransmitter receptors in the STN are needed to infer the various interacting molecular and cellular processes that contribute to these mechanisms. In this study, we constructed a new comprehensive map of connectivity of the inhibitory indirect and excitatory hyperdirect pathways in the mouse and human STN. We identified glutamatergic PV+ STN neurons that show distinct burst firing patterns and topographical segregation in the dorsolateral and middle STN. Cell type-specific synaptic connectivity mapping revealed that both STN PV+ and PV− neurons receive synaptic inputs from both the IP_STN_ and HP_STN_ and that both GPe prototypic and arkypalidal neurons receive synaptic inputs from the HP_GPe_. Thus, “everyone talks, and everyone listens” in cortex-GPe-STN circuitry. These data provide new insights into the basic organization of the BG network and represent an anatomic and functional guide for further dissection of BG functions and DBS effects. Detailed knowledge regarding the anatomy and connectivity pattern of cortico-BG circuits is essential for understanding the psychomotor activity and pathophysiology associated with a wide range of neurological and neuropsychiatric diseases.

The cortex is widely considered to serve as the initial input to the BG through cortico-striatal pathways that convey sensorimotor, associative, and limbic information. The information from these three functional domains is thought to be transmitted to the downstream structures of the BG in a segregated manner, which supports the tripartite model (Lambert et al., 2012; Mallet et al., 2007; Parent and Hazrati, 1995; Plantinga et al., 2018). However, our data indicate that topographically graded organization exists in both the indirect and hyperdirect pathways in the STN, with continuum overlaps, an absence of anatomical boundaries, and direct cortico-pallidal connections. Although previous studies have demonstrated that the coexistence of clearly segregated and overlapping input patterns from the cortex to the BG (Draganski et al., 2008), in the present study we constructed a large-scale map of the indirect and hyperdirect pathways, signifying a new understanding of input organization in the STN. In addition, our detailed analysis of direct cortico-pallidal connections together with recent findings regarding the complexity of the GPe (Abecassis et al., 2020; Mallet et al., 2012; Mastro et al., 2014, 2017) updates current views of the BG network, in which the GPe is considered to be a simple relay nucleus providing a unidirectional flow of information from the input structures (such as the striatum and STN) to the BG output structures (such as the GPi and SNr). These data, together with our single-cell reconstruction and cell type-specific synaptic connectivity analyses, suggest further complexity of BG functional organization through three types of information integration (**Figure 3H**), in addition to reciprocal and loop connections within the BG (Hegeman et al., 2016; Mallet et al., 2012). For instance, beyond the STN-GPe network, the hyperdirect pathway directly and/or collaterally innervates the STN and the striatum as well as two types of GABAergic GPe neurons (prototypic and arkypallidal), thus providing HP-driven feedforward inhibition. This circuit-level complexity implies the existence of anatomical substrates for precisely coordinated regulations of motor as well as nonmotor functions governed by the BG.

It is important to note that, as with any work, the present study has several potential limitations and strengths. First, at the mesoscopic level of mouse connectivity, projection signals labeled using cytosolic fluorescence proteins (FPs) include both axons from passing fibers and synaptic terminals. Like previous studies that found cytoplasmic FP signals to strongly correspond to synaptically-targeted FP signals after removing the axons of passage (Harris et al., 2019; Hunnicutt et al., 2016; Oh et al., 2014), we found no significant difference in the topographic organization of the indirect and hyperdirect pathways after removing the axons of fasciculated passing fibers observed in some areas i.e., the dorsal part of the GPe, and the anterior/posterior ends of the STN (**Figure S4E;** see **STAR Methods**). Second, to test the possibility that our discovery of a topographic gradient in the STN might be the product of graded injections, we examined a dataset showing overlapping axonal projections in the STN that were innervated from separate injections, so that no overlap in these injection sites could be guaranteed. An additional statistical analysis confirmed that neighboring yet non-overlapping injections in different functional subregions led to overlapping projection patterns in the STN (**Figure S4F**). These results support the existence of graded organization in the STN, and challenge the previous models of clear segregation, such as the tripart model. Third, at the microscopic level, although mGRASP-assisted circuit mapping provided convincing data regarding the cell types in the STN and GPe that receive synaptic inputs from the indirect and hyperdirect pathways, further fine-scale mapping of the topographic organization of convergent synaptic connectivity is necessary. Finally, for assessing human connectivity, we used diffusion-weighted image (DWI)-based tractography. This method carries some degree of inherent ambiguity, such as that related to the validity of tract reconstruction and orientation (Maier-Hein et al., 2017; Petersen et al., 2019). However, substantial scientific innovations have aimed to overcome the limitations of this technique. Regardless, it is currently the only tool for mapping human brain connectivity *in vivo,* and the data regarding overlapping patterns were highly consistent with our mouse data, which were obtained using advanced and accurate techniques, as well as with data from previous reports (Plantinga et al., 2016). Overall, our data represent a translational comparison of connectivity patterns in the STN across species.

Previous studies suggested the existence of glutamatergic PV+ neurons in STN and their topographical organization at coarse level (Hontanilla et al., 1998; Roccaro-Waldmeyer et al., 2018; Wu et al., 2018) based on immunohistochemistry with anti-PV or indirect comparison of separate conventional *in situ* hybridization datasets for PV and vGluT2, which makes it difficult to map a precise topographic distribution of STN PV+ neurons. Here, we directly identified topographically localized glutamatergic PV+ neurons in the STN using serial multiplex smFISH. They are a main contributor to burst firing, which indicates a potential key role as generators and modulators of β-band (15-30 Hz) oscillations in the BG. As excess burst firing and β-band oscillations are signatures of Parkinson’s disease and these pathophysiological activities are decreased by STN DBS therapy, these burst firing PV+ STN neurons may be implicated in Parkinson’s disease symptoms (Kühn et al., 2004; Steigerwald et al., 2008). By mapping the distributions of major excitatory and inhibitory transmitter receptors in the STN, we found distinct topographic patterns of expression level. These patterns suggest the existence of a functional layer of differential molecular and cellular receptibility on the top of neural connectivity. Interestingly, the E/I receptor expression ratio is equivalent in dorsolateral PV+ STN neurons, but higher in ventromedial PV-STN neurons. These results raise the important question of whether cell type-specific changes in E/I receptor expression emerge in dopamine-depleted states. Further studies designed to interrogate alterations in synaptic connectivity, receptor expression, and glutamatergic PV+ STN neuronal firing patterns in models of Parkinson’s disease will provide important new insights regarding the coordinated regulation of specialized PV+ neurons in the BG network.

Our systematic connectivity maps, superimposed upon regions with distinct functional cellular heterogeneity, suggest a complex interplay of information within the BG. Our new view of the indirect and hyperdirect pathways (**Figure 7**) substantially updates classical models and proposes, from a structural basis, a new take on relayed and integrated information flow underlying coordinated movement controls and the effects of STN DBS. Further studies of other inputs (e.g. serotonergic and cholinergic) into the STN as well as its outputs will facilitate the elucidation of the mechanisms of the STN functions and complex effects of DBS.

## Acknowledgments

We thank members of the Kim laboratory, cooperation, and technical assistance during the course of the investigation.

## Author Contributions

H.L. performed mouse connectivity experiments and imaging. H.J. performed statistical analysis with mouse connectivity. D.K. and Z.-H.C. provided and analyzed human 7T MRI data. H.L. and Jiwon Kim performed in situ hybridization and immunohistochemistry. K.T.-Y. conducted and analyzed electrophysiological data. H.J., Jiwon Kim and L.F. analyzed molecular profiling data. H.P., Y.L. and S.H.P. performed DBS surgery and collected clinical data. H.J., W.C.O., H.P., and S.H.P. analyzed DBS data. H.L., H.J., D.K., Jiwon Kim., K.T.-Y., H.P., and Jinhyun Kim wrote the original draft. All of the authors reviewed the manuscript. H.L., H.J. and Jinhyun Kim conceived and designed the study.

## STAR Methods

### Mice

C57BL/6J, PV-IRES-Cre (B6;129P2-Pvalb^tm1(cre)Arbr^/J, JAX stock no. 017320), Ai6 (B6.Cg-Gt(ROSA)26Sor^tm6(CAG-ZsGreen1)Hze^/J, JAX stock no. 007906), and double-transgenic PV-IRES-Cre::Ai6 mice were used in this study. While eight 10-week-old mice were used for most experiments, five 7-week-old mice were used for electrophysiology. All experiments were conducted in accordance with protocols approved by the Institutional Animal Care and Use Committee at the Korea Institute of Science and Technology and the National Institutes of Health guidelines for animal care and use.

### Stereotaxic injection

Iontophoresis of recombinant adeno-associated virus (rAAV) vectors was performed by applying a positive current for 5 min using a Digital Midgard Precision Current Source (Stoelting). For indirect pathway mapping experiments, rAAV2/1.CAG.sfGFP.WPRE or rAAV2/1.CAG.tdTomato.WPRE (average titer from seven batches: 1.7 x 10^13^ vg/ml) was injected via a 3 μA current into the GPe. Unilateral dual-injections were performed in 35 mice with a total of 55 injections (**Table S1**). For HP_STN_ labeling, retroAAV-expressing iCre, rAAV2.CAG.iCre.WPRE packaged with a rAAV2-retro helper (Addgene, 51904 and 81070)(Kato et al., 2014; Tervo et al., 2016), was injected via a 5 μA current into the STN of Ai6 mice. For other surgeries, pressure injections were performed at 20–40 nl per min using a Nanoliter injector and a Micro-4 syringe pump (World Precision Instruments). For cortico-pallidal pathway labeling, rAAV2/retro.CAG.iCre.WPRE and rAAV2/1.CAG.JxON.tdTomato.WPRE (100 nl each) were injected into the GPe and ipsilateral primary motor cortex (MOp). For cortico-pallidal pathway dissection, rAAV2/retro.CAG.sfGFP.WPRE and rAAV2/retro.CAG.tdTomato.WPRE (100 nl each) were injected into the STN and the GPe, respectively. For synaptic connectivity mapping, improved mGRASP (Kim et al., 2012) combined with a retro-Lenti system (Kato et al., 2014) or transgenic mouse line was used. For mapping cell type-specific synaptic connectivity, pre- and post-synaptic mGRASP components were injected into the cortex/GPe and STN. mGRASP technology has been demonstrated to be an accurate synaptic detector and has been successfully employed in the mapping of fine-scale synaptic connectivity in multiple cell types in various circuits (Druckmann et al., 2014; Feng et al., 2014; Kim et al., 2012; Kwon et al., 2018; Mukherjee et al., 2020; Song et al., 2018). To validate synaptic connections in hyper-/indirect pathways using mGRASP, rAAV2/1.CAG.pre-mGRASPi.OLLAS.WPRE was injected into the MOp (600 nl) or GPe (400 nl), respectively. For post-mGRASP labeling specific to GPe cell types, 50 nl of retroLenti/FuG-E.CAG.iCre.WPRE was injected into the striatum or STN, and 100 nl of rAAV2/1.CAG.JxON.post-mGRASP.2A.dTomato.WPRE (Addgene, 34913) was injected into the GPe. For post-mGRASP labeling specific to STN cell types, a mixture of rAAV2/1.CAG.post-mGRASP.2A.dTomato.WPRE (Addgene, 34912) and rAAV2/1.CAG.JxON.iRFP670.WPRE (4:1 ratio) was injected into the STN of PV-IRES-Cre mice (50 nl). The following coordinates (in mm) were used for the GPe: −0.22 anterioposterior from bregma (AP), +2.0 mediolateral from bregma (ML), −3.5 dorsoventral from dura (DV); STN: −1.65 AP, +1.6 ML, −4.55 or −4.65 DV (for PV-IRES-Cre); MOp: +2.0 AP, +1.75, +2.0, and +2.25 ML, −0.8 and −0.9 DV; and striatum: +0.5 AP, +2.0 ML, −3.0, −2.5, and −2.0 DV. The viral vectors were packaged using vector cores at the University of Pennsylvania or the University of North Carolina at Chapel Hill.

### Tissue preparation

Brains were prepared as previously described(Feng et al., 2014). For GABA immunofluorescence, 4% paraformaldehyde in 0.1M phosphate buffer with pH 7.4 (4% PFA) containing 0.3% glutaraldehyde was used for fixation. For immunofluorescence, 50-μm sections were permeabilized in 1% Trition X-100 in tris-buffered saline (TBS) and blocked in 5% normal goat serum and 0.4% Trition X-100 in TBS. The sections were incubated with primary antibody for 1–2 days at 4 °C. After washing, the sections were incubated with secondary antibody for 2–3 h at room temperature and counterstained with DAPI, except for in the mGRASP experiments. Sections were mounted with mounting media (Invitrogen, Prolong Diamond Antifade or Vector Labs, VectaShield). In this study, we used the following antibodies: rat anti-OLLAS (Novus, NBP1-06713, 1:1000), guinea pig anti-NeuN (Sigma-Aldrich, ABN90P, 1:1000), rabbit anti-glutamate (Sigma-Aldrich, G6642, 1:500-1000), rabbit anti-GABA (Sigma-Aldrich, A2052, 1:500-1000), Alexa Fluor 488 goat anti-rabbit IgG (Invitrogen, A11008, 1:200), Alexa Fluor 555 goat anti-guinea pig IgG (Invitrogen, A21435, 1:200), Alexa Fluor 633 goat anti-rabbit IgG (Invitrogen, A21071, 1:500), and Alexa Fluor 633 goat anti-rat IgG (Invitrogen, A21094, 1:1000).

For tissue clearing, we modified the CLARITY method (Tomer et al., 2014). Briefly, PFA-fixed brains were cut as parasagittal sections (2.5 mm) containing the GPe and STN. The sections were incubated for at least 2 days in a hydrogel solution with the following composition (in %): 1 acrylamide, 4 PFA, and 0.25 VA-044 initiator in PBS. Following degassing and gel polymerization at 37 °C, the sections were incubated in SmartClear Clearing Buffer Type A (LifeCanvas Technologies) at 37 °C and in EasyIndex (LifeCanvas Technologies) at room temperature.

### Microscopy

Wide-field images were obtained using an Axioscan Z1 slide scanner (Carl Zeiss Microscopy) equipped with a 20X 0.8 NA Plan-Apochromat air lens. Confocal images were acquired using an LSM 780 confocal microscope (Carl Zeiss Microscopy). mGRASP images were obtained at 0.6 μm depth intervals using a 40x 1.4 NA Plan Apochromat oil lens with a 2-fold digital zoom. For cleared tissue imaging, each sample was mounted with EasyIndex. Cleared brain images were acquired at 8.0 μm depth intervals using a 20x 1.0 NA Plan-Neofluar optimized for CLARITY (Carl Zeiss Microscopy). Single-molecule fluorescence *in situ* hybridization (smFISH) images were acquired at 1 or 2 μm depth intervals using a 40x 1.3 NA oil immersion lens.

### Indirect projection mapping pipeline

The MATLAB (https://www.mathworks.com) version of the BaSiC (Peng et al., 2017) algorithm was applied to each fluorescent channel of image tiles (2040 px x 2040 px) using the default parameters for shading correction. Using a custom built GUI, image series were manually inspected for 1) missing or damaged tissue and 2) mispositioning, including flipping and rotation. Image series with severe artifacts in either the GPe or STN were consequently removed from the study. Human experts manually annotated the locations of the injection sites to encompass the cell bodies of labeled neurons, and a morphological operation was performed to generate a smooth binary injection mask. The injection volume of the filtered dataset (n = 55) averaged 0.1403 ± 0.0797 mm^3^ (0.0546 ± 0.031 mm^3^ within the GPe).

The custom signal detection algorithm was applied to each image to extract positive signals from the background autofluorescence. The autofluorescence distribution of each image series was estimated from the non-signal regions contralateral to the injection site, and the mean was subsequently subtracted from the image. A median filter (3 x 3) was applied to reduce the noise. Candidate signal objects were first detected using the autofluorescence intensity profile and classified into three groups based on the intensity. Low- and medium-intensity objects were combined with the detection result from the edge detection filter (k_width_ = 10, 30). The binary signal masks generated from the detection results were used to classify each pixel as either signal or background, and the density within a 10 μm × 10 μ grid was computed.

Images were first pre-aligned sequentially to minimize the inter-sectional difference in the DAPI signal. Stacked image volumes were aligned to the Allen Common Coordinate Framework (CCF v3) (Wang et al., 2020) via iterative 3D global and two-dimensional (2D) slice affine alignment using elastix (Klein et al., 2010). A population average model from 35 aligned volumes was generated following the Allen informatics data pipeline (Kuan et al., 2015), then registered to the CCF v3 using the Symmetric Normalization (SyN) transformation model (Avants et al., 2008) from Advanced Neuroimaging Tools (ANTs, https://github.com/ANTsX/ANTs). 3D deformable transformation from the population-averaged model to the CCF v3 was applied to each DAPI-based volume and the corresponding signal image was used to generate a 25 μm isotropic 3D registered volume. Injection and projection density signals within the GPe and STN were segmented using 3D region annotation from the Allen Mouse Brain Atlas and stored in a vectorized form. Signals in the GPe were downsampled to an isotropic voxel-size of 50 μ facilitate computation. Signal volumes were computed as the sum of the signal density within the target structure, multiplied by the voxel size.

To limit false positive projection signals, image series with (1) low injection or projection volume (< 0.001 mm^3^) or (2) ectopic injections at the central nucleus of the amygdala (CeA) or substantia innominata (SI) were removed.

### Hyperdirect projection data from the AMBCA

Using the API for the Allen Mouse Brain Connectivity Atlas (AMBCA, https://connectivity.brain-map.org), we downloaded the dataset of isocortical injections with GPe or STN projections (n_gpe_ = 1270, n_stn_ = 851, at the time of our analysis). The voxel resolution for isotropic volume was set to 50 μm for injections and the GPe projection density, and 25 μm for the STN projection density. Injection sites were inspected to remove experiments in which more than 10% of the injection volume was located outside the isocortex. The minimum projection volume for both the GPe and STN was set to 0.001 mm^3^. We included nine transgenic Cre-lines with at least five datasets (Table S3) in addition to a dataset for wild-type C57BL/6J, which result in 176 datasets in total (181 for HP_STN_).

Trainable WEKA Segmentation (http://fiji.sc/Trainable_Weka_Segmentation) plugin for Fiji was used to identify and subtract passing fibers in the GPe and STN. The WEKA Segmentation model was trained using set of manually annotated labeled images containing labels for 1) densely populated thick fasciculated axon fibers, 2) fine axon projections, and 3) background. Among 86 datasets with densely populated projection signal (density > 0.5) in GPe, 25% were randomly selected for training dataset (*n_train_* = 22). Selection of filters and parameters were same as described in Hunnicutt *et al*(Hunnicutt et al., 2016). After training, the classifier was applied to remaining dataset and generated a probability map for each label. To minimize the artifacts from thick fasciculate fibers, conservative segmentation criteria was set to subtract as much passing fiber signal as possible. Default classification threshold was set to *P_fiber_* > 0.25 for fasciculated fiber, and *P_signal_* > 0.7 for axon signal classification. Custom built GUI in MATLAB was used to inspect the classification result, and to manually tune the segmentation threshold for each data if needed.

### Single-neuron reconstruction data from the MouseLight database

Single neuron reconstruction data were obtained from the MouseLight database using Neuron Browser (http://www.mouselight.janelia.org). The search query was set to include only neurons for which the soma was located within the isocortex and any axonal structures were in either the GPe or STN (**Figure S5B**), which resulted in 142 reconstructed neurons. Neurons whose soma was located in the left hemisphere were mirrored onto the right hemisphere. Reconstructed neurons without ipsi-lateral projections to the GPe or STN were removed, and the remaining neurons (n = 69) were classified by projection pattern: (1) IT types showing contralateral projections (n = 24/69); (2) Collateral HP_STN/GPe_ (n = 10/69); (3) HP_STN_ only (n = 15/69); and (4) HP_GPe_ only (n = 20/69).

Normalized projection density was computed as the length of axons within the region normalized by the region volume. To visualize the topographical organization (Figure S5D), only neurons with axonal projections longer than 100 μm in both the GPe and STN were shown.

### Analysis of mouse hyper/indirect pathway topographic organization

The projection centroid was defined as the weighted centroid of the 3D density signal within a target region. The distance between two projections was measured using the pairwise cosine distance between two vectorized projections. STN geometric axis1 and axis2 were defined as the principal axis and its perpendicular, respectively, in a 2D projection of the STN along the AP axis.

For the signal distribution heatmap, reconstructed signals were digitally resliced along a designated axis and the fraction of signal volume within each slice to the total signal volume was computed. The similarities between signal distributions were measured using the Pearson correlation between the signal centroid location along each direction.

We applied classical multi-dimensional scaling (MDS) (Kruskal and Wish, 1978) to visualize the pairwise projection distance relationships between STN projections in the “pattern space”. Robust Canonical Correlation Analysis (rCCA) (Branco et al., 2005) was performed to identify the graded topographic organization between the locations of each projection in the pattern space and the injection centroid in the anatomical space. MDS embeddings were then rotated so that the major axes corresponded to the canonical variates. Gradient axes in the cortex and the GPe corresponding to the STN geometric axes were computed via ridge regression on the STN projection centroid location along the gradient axis and input centroid coordinates.

Injections for which the majority (> 80%) of the injection volume was contained within the primary injection structure were classified as localized injections and used to investigate projection patterns from a single cortical subregion. Projection density signals from localized injections into the target structure were rescaled to [0, 1], then averaged to generate a characteristic projection. Note that the resulting characteristic projection may reflect only the spatial pattern of the projection, and its intensity may not be comparable to characteristic projections from other regions.

The projection power of a functional subregion, which represents an estimated projection volume assuming the subregion is fully labeled, was computed using non-negative ridge regression on the projection volume with the vector containing the injection volume within each subregion normalized by its volume, as a regressor.

The topographic relationship between IP_STN_-IP_INPUT_ was modeled using a multivariate linear model with the IP_STN_ centroid locations along two gradient axes as observations and the IP_INPUT_ injection centroids as predictors. The same model was assessed using HP_STN_ centroids as inputs and HP_GPe_ centroids as targets. Error was measured according to the distance between the actual and predicted HP_GPe_ centroids. Permutation tests were used to compare the means of observed error against the error from 10,000 randomizations, where for each HP_STN_, the corresponding HP_GPe_ was reassigned by shuffling. Two-sided t-tests were used to compare the means of the two groups.

In addition, to test whether nearby cortical injections targeting different functional subregions resulted in clearly segregated STN projection patterns, we measured the STN projection pattern distances between pairs of experiments using the following criteria: 1) the two injection volumes are localized within different functional tripartite cortical subregions; 2) the injection volumes exhibit no overlap in the reference space; 3) the distance between injection centroids is greater than 1 mm and less than 1.5 mm. We compared the distribution and mean of the STN projection distances from pairs of nearby injections targeting different subregions and pairs with the same input subregion using two-sided t-tests.

### Serial Single-Molecule Fluorescence In Situ Hybridization

Mouse brains were collected and immediately frozen in −80 °C 100% ethanol. Coronal sections (20 μm) containing the STN were collected using a cryostat (Leica) and mounted on SuperFrost microscope slides (FisherBrand). Serial smFISH was performed using the RNAscope multiplex fluorescent kit (ACD bio, 320850) according to the manufacturer’s protocols. Brain sections were fixed in 4% PFA and dehydrated serially in ethanol (50%, 70%, 100%, and 100%). Sections were incubated in proteinase IV and rinsed with PBS. Probes were applied for 2 h at 40 °C and amplifiers 1–4 were subsequently applied at 40 °C. We used probes for *Aqp4* (catalog no. 417161), *Gabra1* (435451), *Gria1* (421931), *Mbp* (451491), *Pvalb* (421931), *Rbfox3* (313311), *Slc17a6* (319171), *Slc32a1* (319191), and *Tmem119* (472901). Sections were counterstained with DAPI.

A nucleus detection pipeline, as previously described (Feng et al., 2019), was trained on 30 images containing DAPI-stained cell nuclei (∼180 per image) that were manually labeled by human experts. To detect mRNA puncta, we used a splitting strategy with a marker-controlled watershed approach and a variational Bayesian Gaussian mixture model (Feng et al., 2012). The detected mRNA puncta were assigned to the nearest detected nuclei with a maximum distance of 1 μm.

Human experts manually registered each smFISH image section to a corresponding slice in Allen CCF v3, using a GUI that was custom designed in MATLAB, via rigid transformation (reflections, translation, and rotations). Support Vector Machine was then used to determine the threshold for positive cell classification. A classification model was trained on 30 randomly selected and manually labeled cells per image section, using the number, size, and intensity of puncta as the input.

### Electrophysiological recording

Mice were anesthetized with halothane and transcardially perfused with an ice-cold dissection solution containing (in mM): 228 Sucrose, 2.5 KCl, 7 MgSO_4_, 0.5 CaCl_2_, 1.25 NaH_2_PO_4_, 26 NaHCO_3_, and 11 glucose, oxygenated with 95% O_2_ and 5% CO_2_. The brains were quickly removed and parasagittal slices (250 μm) containing the STN were prepared using a vibratome (Leica VT 1200S) in cold dissection solution. Slices were then transferred into extracellular solution containing (in mM): 125 NaCl, 2.5 KCl, 1.3 MgSO_4_, 2.5 CaCl_2_, 1.25 NaH_2_PO_4_, 26 NaHCO_3_, and 11 glucose, oxygenated with 95% O_2_ and 5% CO_2_, in which they were constantly kept before being moved to the recording chamber which also contained the extracellular solution. Chemicals for electrophysiological analyses were obtained from Sigma or Wako Pure Chemical Industries unless otherwise specified.

Whole-cell patch clamp recordings were made from ZsGreen-positive or ZsGreen-negative neurons in the STN of PV-IRES-Cre::Ai6 mice. Patch pipettes (resistance 5-6 M Ω) were filled with a solution containing (in mM): 130 potassium gluconate, 2 NaCl, 4 MgCl_2_, 4 Na_2_ATP, 0.4 NaGTP, 20 HEPES (pH 7.2), and 0.25 EGTA. Recordings in the cell-attached configuration were sometimes conducted before making whole-cell recordings. Neurons in the STN were visualized directly using a BX61WI microscope (Olympus), and those expressing ZsGreen were identified by a FV1000 confocal microscope (Olympus). Membrane potentials were recorded in current-clamp mode either without current injection, or by giving depolarizing (100 pA) or hyperpolarizing (−200 pA) current steps for 500 ms, using a Multiclamp 700B amplifier and pClamp software (Molecular Devices). Data were accepted if the input membrane resistance was > 100 M Ω, firing action potentials had an amplitude of at least 50 mV, and the afterhyperpolarizing potential was < −50 mV. Analyses of electrophysiological parameters were performed using OriginPro (https://www.originlab.com) and Clampex (https://www.moleculardevices.com) software. The peak of afterhyperpolarizing potential was considered as membrane potential during firing. The parameter values for action potentials were equivalent to a previous study (Beurrier et al., 1999). Statistical significance was evaluated using the Mann-Whitney U test. In this study, to examine two neuronal firing patterns (tonic and burst firing), we classified burst firing according to the presence of, at least, one of the following firing patterns in three sequential recording conditions: 1) When recording without current injection, spontaneous firing shows fast burst firing (< 10 ms inter-spike intervals, ISI, on average) followed by a pause. Similar to the previously reported burst firing in the STN (Beurrier et al., 1999; Huang et al., 2017; Kass and Mintz, 2006), the membrane potential tends to be depolarized during burst firing, and tends to be drastically hyperpolarized at the start of a pause; 2) when recording during a 250-ms period of rebound depolarization following the removal of a hyperpolarizing current injection (−200 pA), the firing frequency is fast (< 10 ms ISI on average) and the membrane potential tends towards depolarization, followed by drastic hyperpolarization (> 15 pA) and quiescence of firing; 3) when recording during a 500-ms depolarizing current injection (100 pA), the firing frequency is fast (< 10 ms ISI on average) and the membrane potential tends towards depolarization at the beginning of a depolarizing current injection, followed by quiescence of firing for more than 50 ms (e.g., the top-right trace in **Figure S9C**). The firing properties used for this classification were compared between burst PV+, tonic PV+, and tonic PV-neurons. We noted that the properties of burst firing neurons generally differed from those of tonic firing neurons and that there was a clear tendency for firing to slow down during bursts, as reflected by the significant difference between the first or second ISI and the last ISI.

### Human 7T-DTI tractography

T2*-weighted imaging (T2*WI) with 0.2 mm in-plane resolution and diffusion-weighted imaging (DWI) along 64 directions with a b-value of 2000 s/mm^2^ and b0 with 1.8 mm isotropic resolution were obtained from a healthy 30-year-old adult male using a 7.0T MRI scanner (Magnetom, Siemens) and a homemade 8-channel radiofrequency head coil (Gachon University) (Cho et al., 2014). A 3D T1-weighted whole brain scan was obtained for anatomical reference using a 3.0T MRI scanner (Magnetom Verio, Siemens).

The DWI data were preprocessed to correct geometric distortions (Oh et al., 2012). Axonal fiber orientation information was calculated using a constrained spherical deconvolution-based diffusion model in the MRtrix3 toolbox (https://www.mrtrix.org)(Tournier et al., 2010). Whole-brain streamline tractography was performed using the probabilistic algorithm, iFOD1, as implemented in the MRtrix3 and visualized by track-density imaging (TDI) (Calamante et al., 2010) with a 0.2 mm isotropic resolution. Stacked T2*-WIs were interpolated to an isotropic 0.2 mm resolution and then co-registered to the b0 volume image of the DWI data using SPM12 (https://www.fil.ion.ucl.ac.uk/spm/software). The regions-of-interest, including the STN and GPe, were manually traced using the MRtrix3, and the 3D Slicer (http://www.slicer.org) was used for 3D modeling. T1-images were registered to the b0 volume image of DWI scans via affine transformation and annotated using the human Brainnetome atlas (Fan et al., 2016), mapped using FreeSurfer (http://surfer.nmr.mgh.harvard.edu).

Probabilistic constrained spherical deconvolution-based fiber tracking was performed using the iFOD2 algorithm (Oh et al., 2012). Optimal parameters were determined such that the resulting streamlines were visually similar to the combined subset of streamlines filtered according to anatomical constraints (Desmond et al., 1995). An additional five parameter combinations with values close to the selected parameters were also used. Whole-track and end-point TDI maps were computed using tracks that terminated within the STN.

The normalized average density of tracks in a region was defined as the average track intensity of all voxels within the TDI map of the target subregion. The spatial overlap was defined as 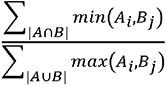, where A_j_ and B_j_ are the voxel intensities from two subregions.

### Human deep brain stimulation

This study included 26 patients who received bilateral STN DBS implantation between March 2005 and December 2008 at the Movement Disorder Center of Seoul National University Hospital (SNUH). The criteria were electrical stimulation of the left STN with a mono-polar electrode configuration and follow-up period of more than 3 years.

Pre-operative MRI images were obtained using a 1.5 T Signa MRI system (General Electric Medical Systems). STN DBS implantation was performed using a Leksell frame and SurgiPlan with microelectrode recording and macro-stimulation under local anesthesia, as previously described (Kim et al., 2009, 2010). The dorsolateral portion of the STN was targeted using MRI images, with theoretical STN coordinates (4 mm anterior from the midcommissural point, 10–12 mm lateral from the midline, and 3–4 mm below the intercommissural line). Permanent DBS electrode (Medtronic, DBS 3389) placement was determined to avoid damage to adjacent vessels, ventricles, and sulci.

Three-dimensional spiral stereotactic CT scans (Philips, Brilliance CT 64-channel) with 1 mm thick slices were taken more than 1 month after surgery as previously described(Paek et al., 2008). The post-operative CT images were co-registered to pre-operative MRI images that were normalized to the ICBM 152 2009b nonlinear asymmetric T2 template (Fonov et al., 2011). The electrode trajectories were pre-reconstructed from post-operative CT images and manually adjusted. All normalization and registration processes were done using Lead-DBS (Horn and Kühn, 2015). Electrode trajectories were plotted onto images from the Schaltenbrand and Wahren human brain atlas for 2D visualization (Schaltenbrand and Wahren, 1998).

Neurologists evaluated the patients using the Unified Parkinson Disease Rating Scale (UPDRS) before and at 12, 24, 36, and 60 (if applicable) months after surgery. Motor symptom improvement was measured as a post-operative change in the summed UPDRS motor score (UPDRS III) of the right limb.

### Data Availability

The datasets generated and analyzed supporting the findings of this study are available from the corresponding author upon reasonable request.

### Code availability

Custom code used in this study are available from the corresponding author upon request.

### Supplemental Information

**9** Figures; **6** Tables; **2** Movies

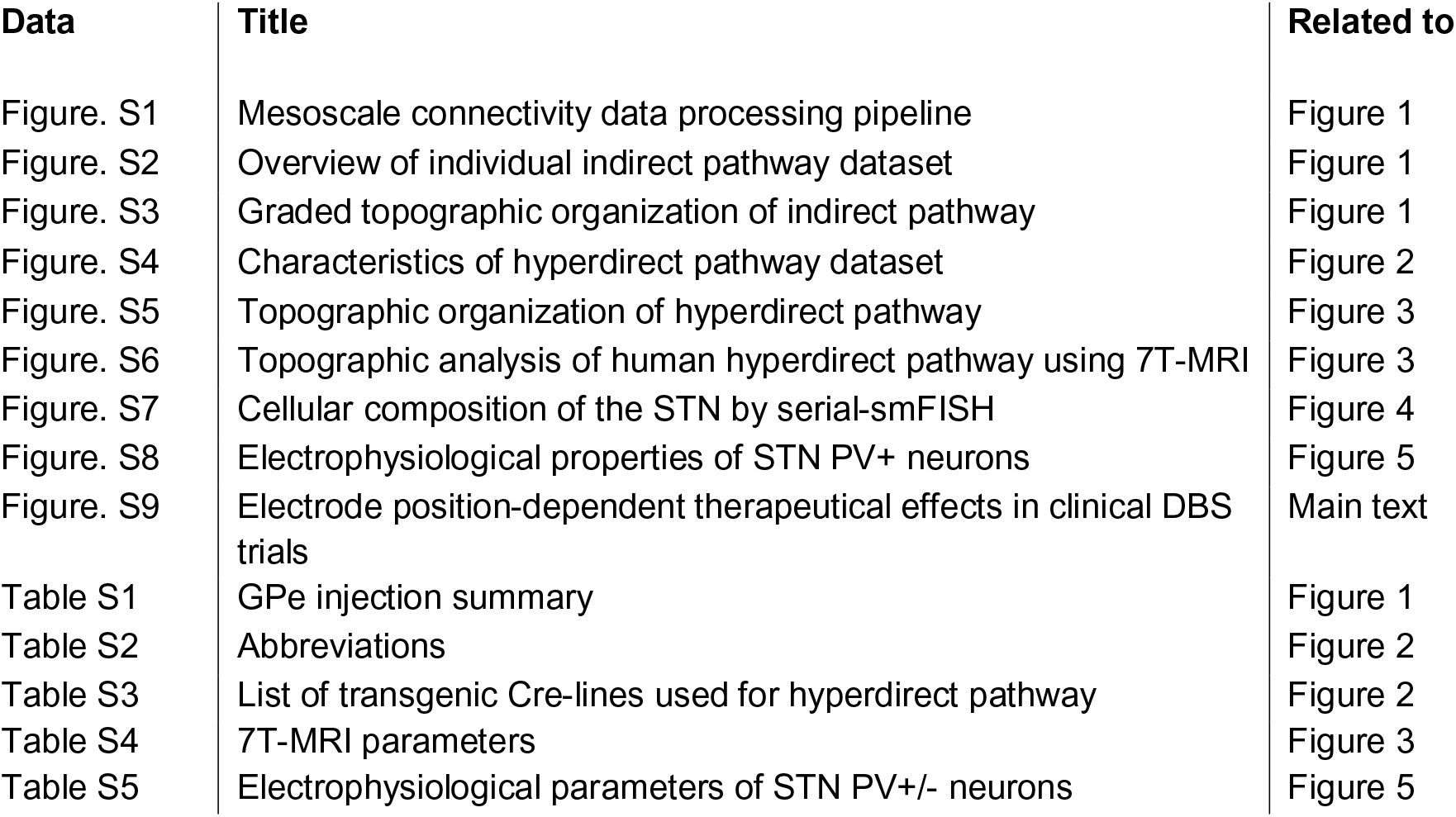
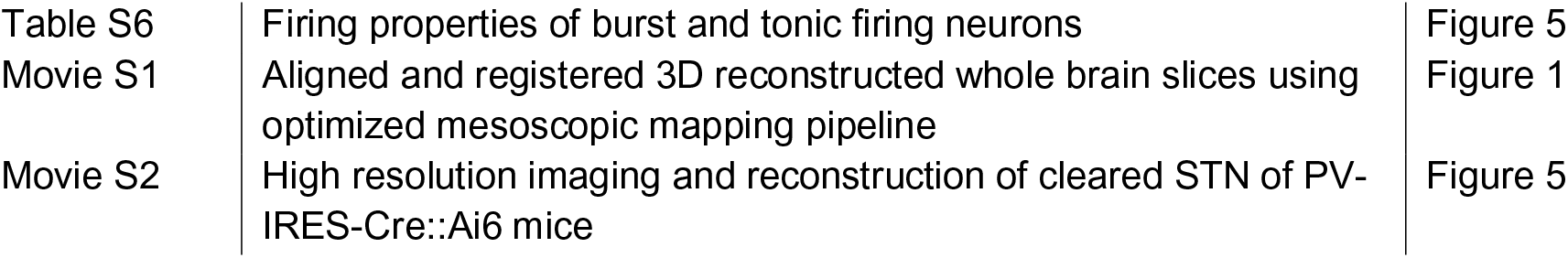

