## Supplemental Information for "Topographic connectivity and cellular profiling reveal detailed input pathways and functionally distinct cell types in the subthalamic nucleus"

**Supplemental Information**
**9** Figures; **6** Tables; **2** Movies

| **Data** | **Title** | **Related to** |
|---|---|---|
| Figure. S1 | Mesoscale connectivity data processing pipeline | Figure 1 |
| Figure. S2 | Overview of individual indirect pathway dataset | Figure 1 |
| Figure. S3 | Graded topographic organization of indirect pathway | Figure 1 |
| Figure. S4 | Characteristics of hyperdirect pathway dataset | Figure 2 |
| Figure. S5 | Topographic organization of hyperdirect pathway | Figure 3 |
| Figure. S6 | Topographic analysis of human hyperdirect pathway using 7T-MRI | Figure 3 |
| Figure. S7 | Cellular composition of the STN by serial-smFISH | Figure 4 |
| Figure. S8 | Electrophysiological properties of STN PV+ neurons | Figure 5 |
| Figure. S9 | Electrode position-dependent therapeutical effects in clinical DBS trials | Main Text |
| Table S1 | GPe injection summary | Figure 1 |
| Table S2 | Abbreviations | Figure 2 |
| Table S3 | List of transgenic Cre-lines used for hyperdirect pathway | Figure 2 |
| Table S4 | 7T-MRI parameters | Figure 3 |
| Table S5 | Electrophysiological parameters of STN PV+/- neurons | Figure 5 |
| Table S6 | Firing properties of burst and tonic firing neurons | Figure 5 |
| Movie S1 | Aligned and registered 3D reconstructed whole brain slices using optimized mesoscopic mapping pipeline | Figure 1 |
| Movie S2 | High resolution imaging and reconstruction of cleared STN of PV-IRES-Cre::Ai6 mice | Figure 5 |

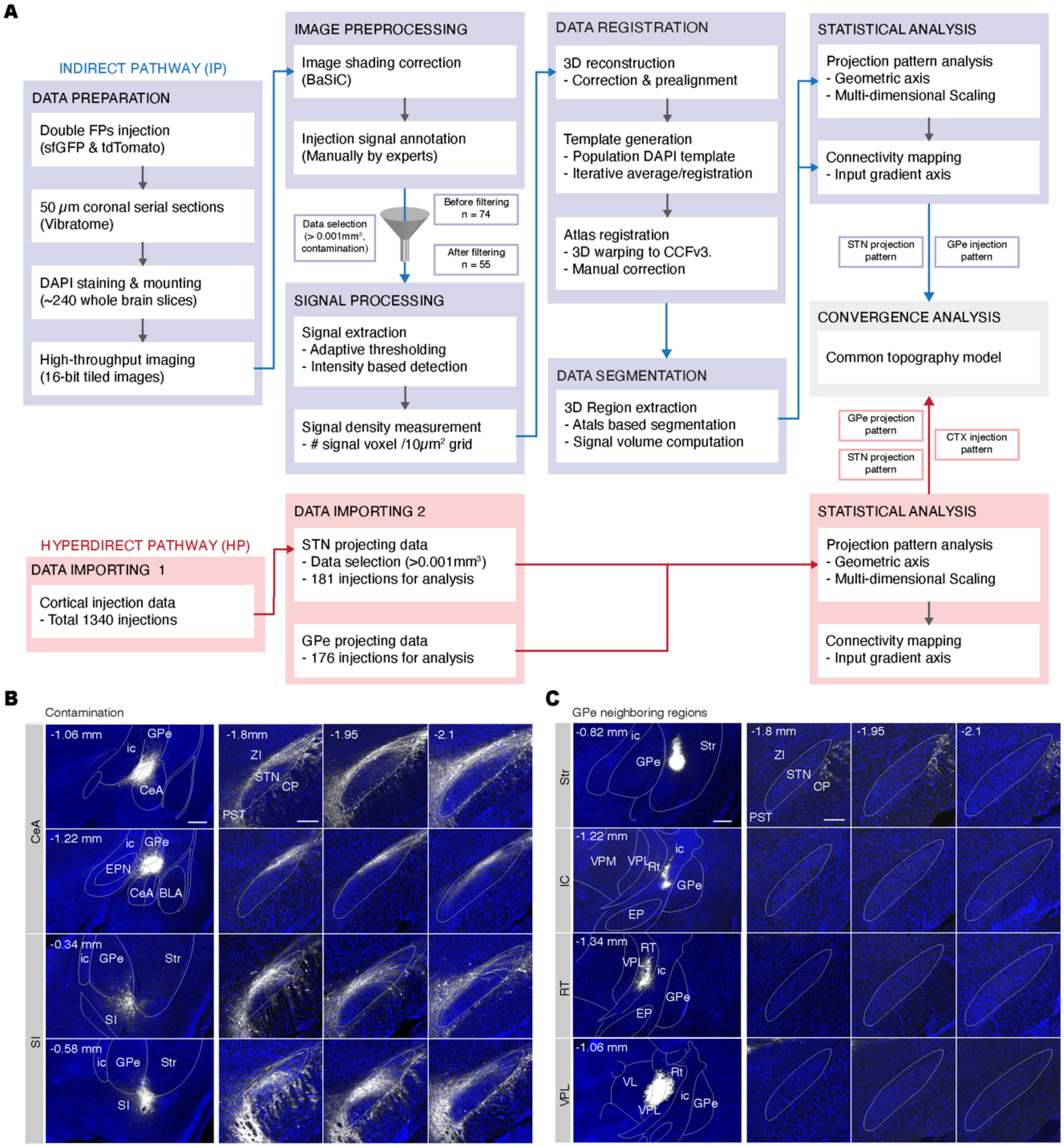


**Figure S1. Mesoscale connectivity data processing pipeline, Related to Figure 1. (A)** Workflow of data generation and analysis for the indirect pathway obtained by GPe injections (blue flow chart) and the hyperdirect pathway using the Allen Mouse Brain Connectivity Atlas (AMBCA) (red flow chart). **(B), (C)** Ectopic injections in neighboring regions around the GPe with **(B)** and without **(C)** axonal projections in the STN, which were excluded for further indirect pathway analysis. Coronal images of the GPe at the center of the injection sites (left panel) and the STN at three points along the AP axis (right), with numbers indicating the distance from bregma. Scale bar, 250 μm (left); 100 μm (right).

**A**

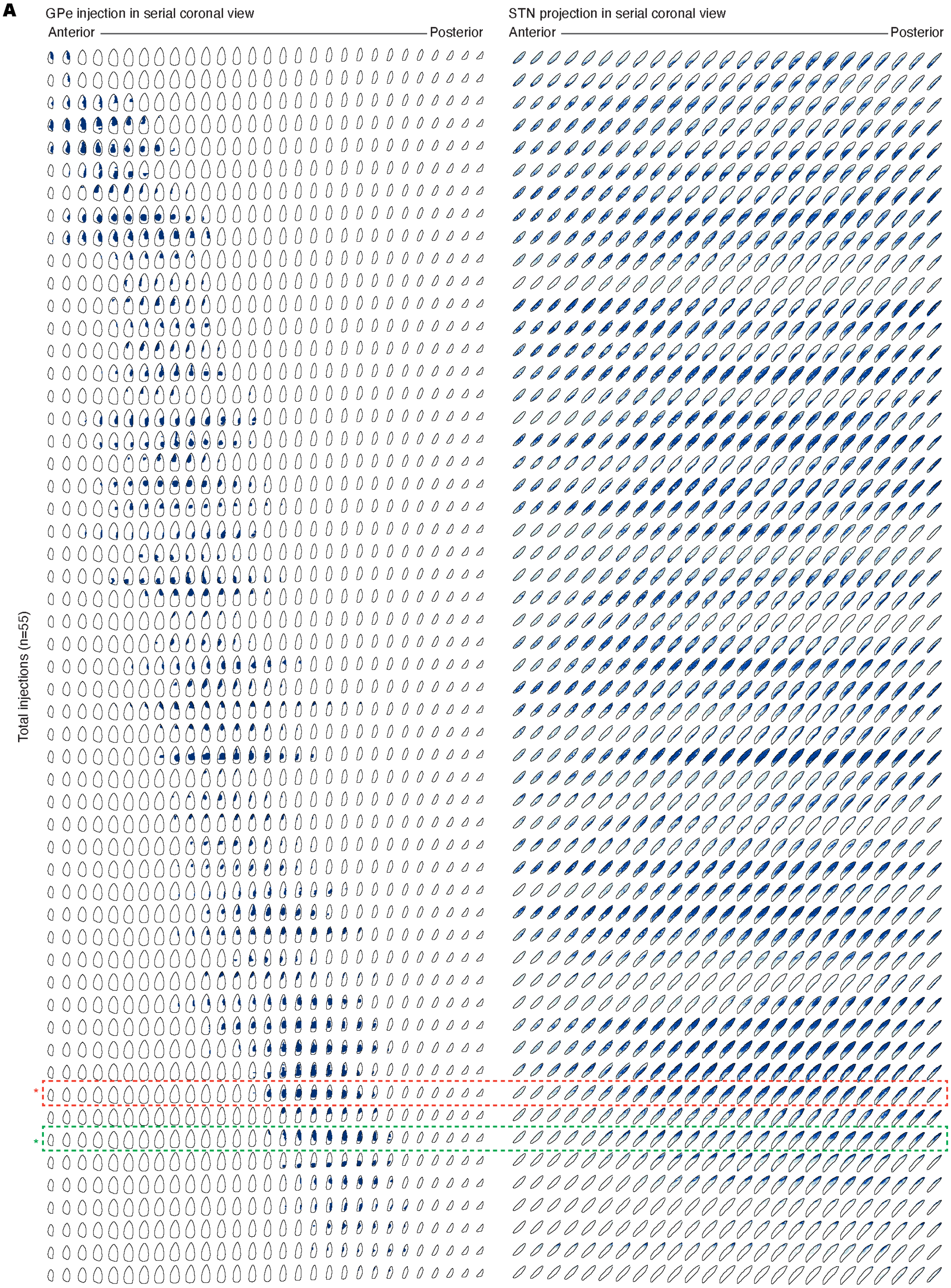


**Figure S2. Overview of the individual indirect pathway dataset, Related to Figure 1.** Digitally reconstructed serial coronal view of all injections, indirect pathway input (left, n = 55), and the corresponding projections, and $IP_{STN}$ (right), presented side-by-side (AP interval of 50 μm for the

GPe; 25 µm for the STN). Injections are shown using binary masks annotated by human experts, and the projections are shown in terms of projection density. Rows are sorted according to the AP coordinates of the injection centroid. Experiments denoted by the red and green box indicate the example datasets shown in **Figure 1G (top)**.

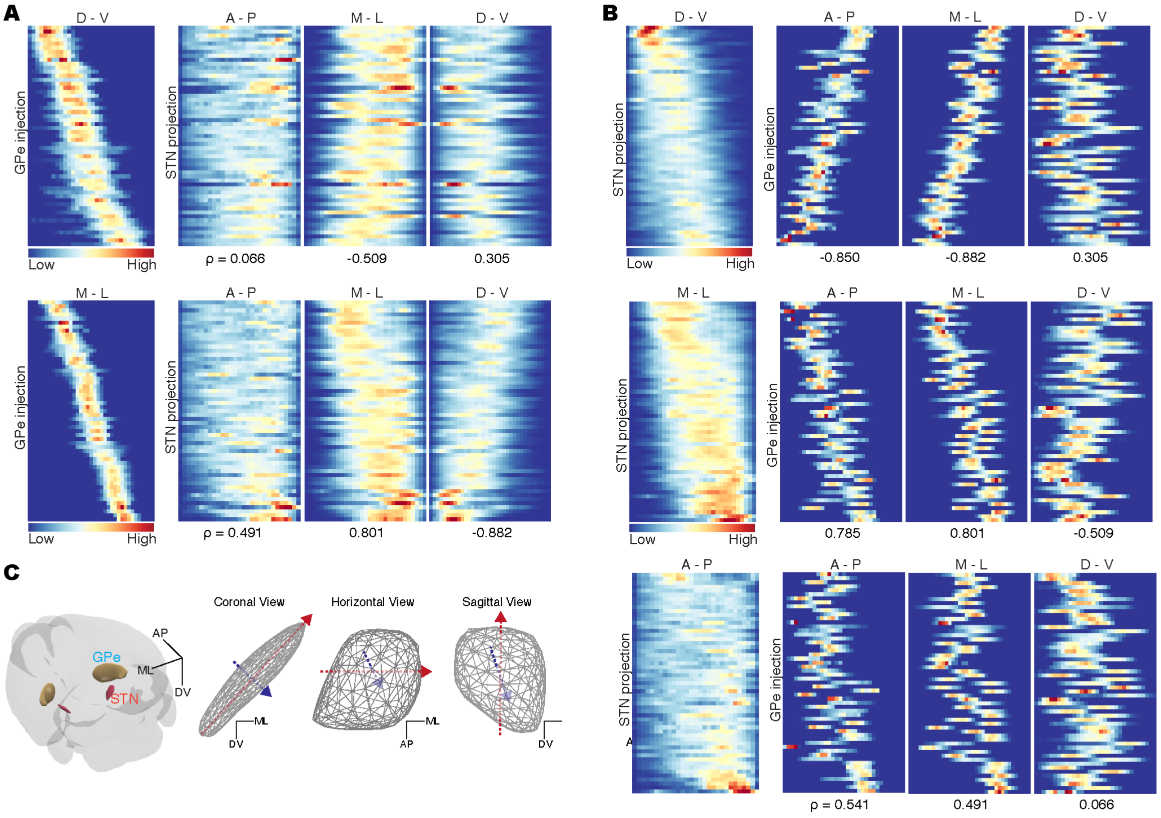


**Figure S3. Graded topographic organization of the indirect pathway, Related to Figure 1. (A), (B)** Heatmap of the indirect pathway input, and $IP_{STN}$ signal distribution along anatomical axes (AP, anterior-posterior; ML, medial-lateral; and DV, dorsal-ventral). Data are sorted according to GPe injection **(A)**, and $IP_{STN}$ **(B)** centroid locations along each anatomical axis. Correlations between indirect pathway input and $IP_{STN}$ were measured using the Pearson's r.

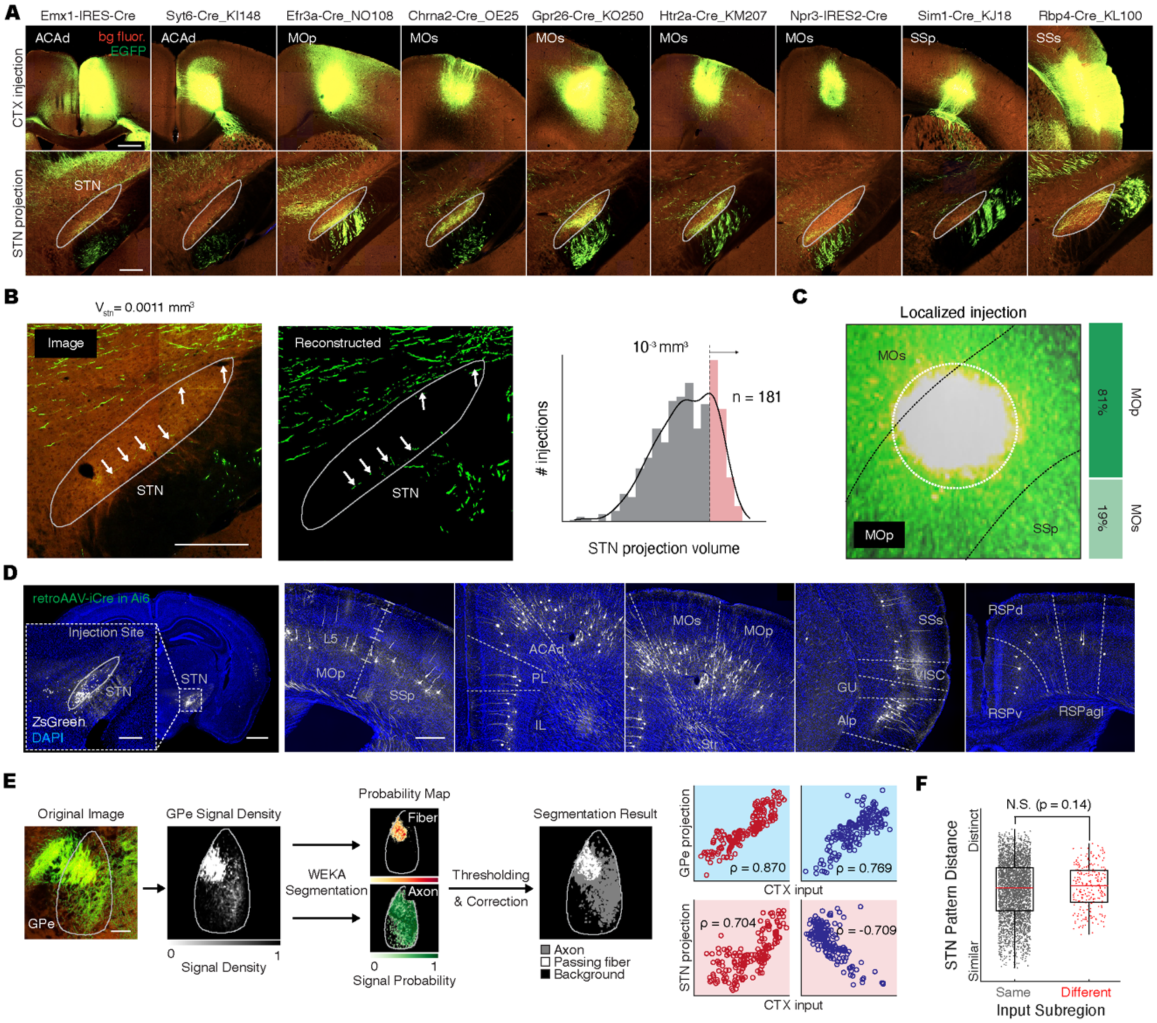


**Figure S4. Characteristics of hyperdirect pathway dataset, Related to Figure 2. (A)** Example images of cortical injections in 9 transgenic mouse lines showing axonal projections in the STN, extracted from AMBCA datasets. Scale bar, 500 μm (top); 250 μm (bottom). **(B)** Left: example of an STN axonal projection image with low signal (just above the threshold, $10^{-3}$ $mm^3$) and corresponding segmented signal (left). White arrows indicate thin axons within the STN. Scale bar, 250 μm. Right: the distribution of log10-transformed STN projection volumes from 851 cortical injection datasets with fitted non-parametric kernel-smoothing distribution. The distribution of the STN projection volume for the 181 selected injections is shown in red. **(C)** Example of a cortical injection image of a localized injection in the MOp (AMBCA, 298273313) illustrating the threshold criteria for data filtering (> 80% injection volume in a single subregion), which was used for projection description of an individual subregion. **(D)** Injection site of retroAAV-iCre in the STN of ZsGreen reporter mice (Ai6) (left) and an example of labeled STN projecting cells across the cortical subregions. Scale bar, 1 mm; 250 μm (inset). **(E)** Left: Workflow for segmenting fasciculated passing fibers using Advanced WEKA Image Segmentation and example result of processed GPe projection image (AMBCA, 100141780). Scale bar, 250 μm. Right: linear relationship between axial locations of the $HP_{GPe}$ (top), $HP_{STN}$ (bottom), and cortical inputs after removing passing fibers, computed using the same analysis as in **Figure 3D. (F)** Distribution of STN projection pattern distance for neighboring yet non-overlapping injections, from the same vs. different functional subregion.

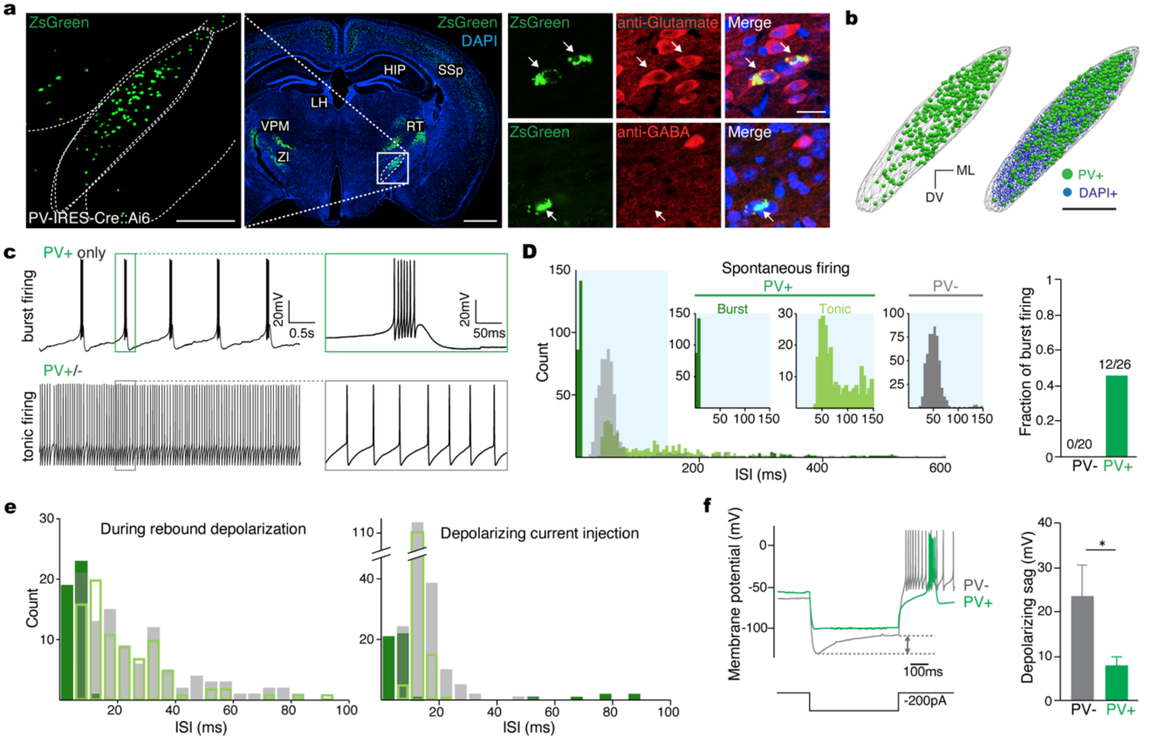


**Figure S5. Topographic organization of hyperdirect pathway, Related to Figure 3. (A)** Heatmap of sorted hyperdirect pathway input with $HP_{STN}$ and $HP_{GPe}$ signal distribution along anatomical axes. The functional parts of each cortical injection (row) are color-coded (somatomotor, limbic, and association in green, red, and yellow, respectively). Rows are sorted by the AP, ML, and DV location of injection centroid, respectively. **(B)** Top: example of a search query used to download all neurons showing either $HP_{GPe}$ or $HP_{STN}$ projections from the MouseLight database. Bottom: heatmap of axonal projection density within basal ganglia structures from reconstructed cortical neurons showing ipsilateral STN or GPe projections. IT/PT cell types were classified based on the existence of contralateral projections (data not shown), and collateral projection neurons were marked by a dashed box. **(C)** 3D visualization of 45 PT-type neurons with axonal ends on either the GPe or STN in sagittal (left) and horizontal views (middle), see enlarged views in the dashed boxes (right). (**D**) Left, soma locations of 9 collateral $HP_{GPe/STN}$ projection neurons whose axonal projections to both GPe and STN were longer than 100 μm (left). Axonal projection of collateral $HP_{GPe/STN}$ projection neurons to the GPe (right, top) and STN (right, bottom). Centroid of soma, shown in dots, and axonal projections are colored based on soma location along CTX gradient axis1. Arrows represent the STN geometric axis and corresponding GPe gradient axis.

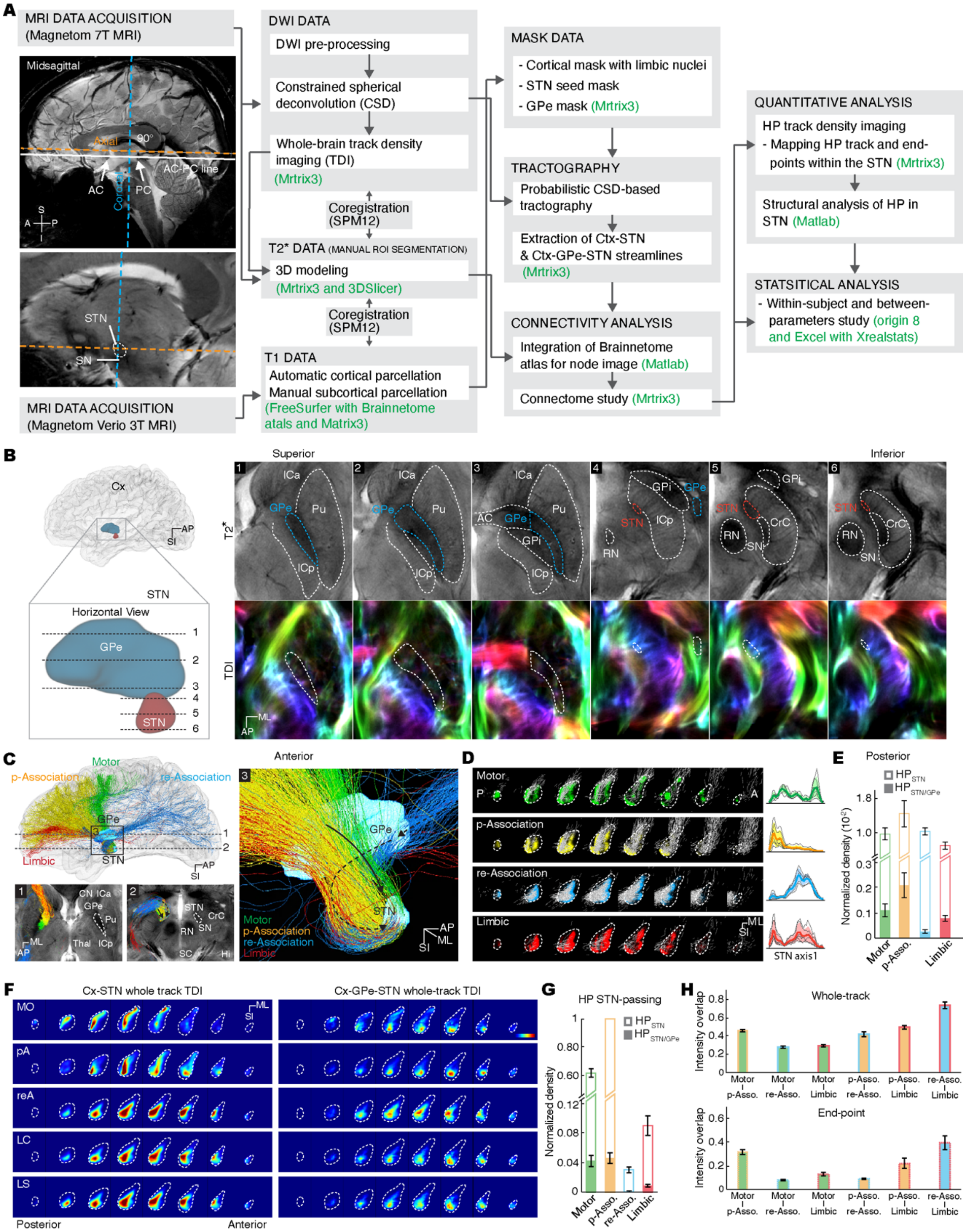


**Figure S6. Topographic analysis of human hyperdirect pathway using 7T-MRI, Related to Figure 3. (A)** Pipeline of 7T MRI and tractography data acquisition and processing. Midsagittal MR images illustrate the imaging reference line set along the central intercommissural line. Software tools used for processing are specified within each step (also see **STAR Methods** for details of data processing, **Table S4** for scan parameters). **(B)** 7T T2*-weighted images of the GPe and STN in axial views. 3D representation of the cortex, GPe, and STN, with imaging locations shown from a midsagittal view (left). Anatomical annotations drawn by a human expert (right, top) and directionally-encoded color track density images shown on T2*-weighted images (right, bottom). Fiber-directionality information was incorporated as red (ML)–green (AP)–blue (SI). **(C)** 7T MRI-based tractography of the human $HP_{GPe/STN}$ in midsagittal 3D view (left top). 2 mm thick axial T2*-weighted image overlaid with streamline tracks of the $HP_{GPe/STN}$ (1, GPe level; 2, STN level, bottom) and enlarged $HP_{GPe/STN}$ streamline tracks in 3D (3, right). Tracks are colored according to connections with cortical functional parts (motor, limbic, prefrontal (p)- and rest (re)-association). Arrows indicate two distinct HP trajectories. AP; anterior-posterior, ML; medial-lateral, SI; superior-inferior. **(D)** Serial coronal (0.65 mm spacing) STN end-point track density imaging (TDI) with reconstructed whole-track $HP_{STN}$ streamline in white (left). Line plot of whole-track TDI signal distribution along STN axis1 (right). Data are shown as means (bold) with SD, with individual tracking results indicated by thin gray lines (n = 6). **(E)** Normalized average density of $HP_{GPe/STN}$ and $HP_{STN}$ end-point tracks. Error bars denote s.e.m. **(F)** Serial coronal probabilistic tractography-based whole-track density image (TDI) of the $HP_{STN}$ (left) and $HP_{GPe/STN}$ (right). Pseudocolors represent track density. **(G)** Normalized average density of tracks including STN-passing projections for the $HP_{GPe/STN}$ (colored bar) and $HP_{STN}$ (outline). Density values were normalized to the maximum average density ($HP_{STN}$ from the prefrontal association cortex). **(H)** Bar plot showing the whole-track density (top) and end-point density (bottom) overlap within the STN between $HP_{STN}$ connected to each cortical functional part. The color and edge of each bar represent a compared pair of functional parts.

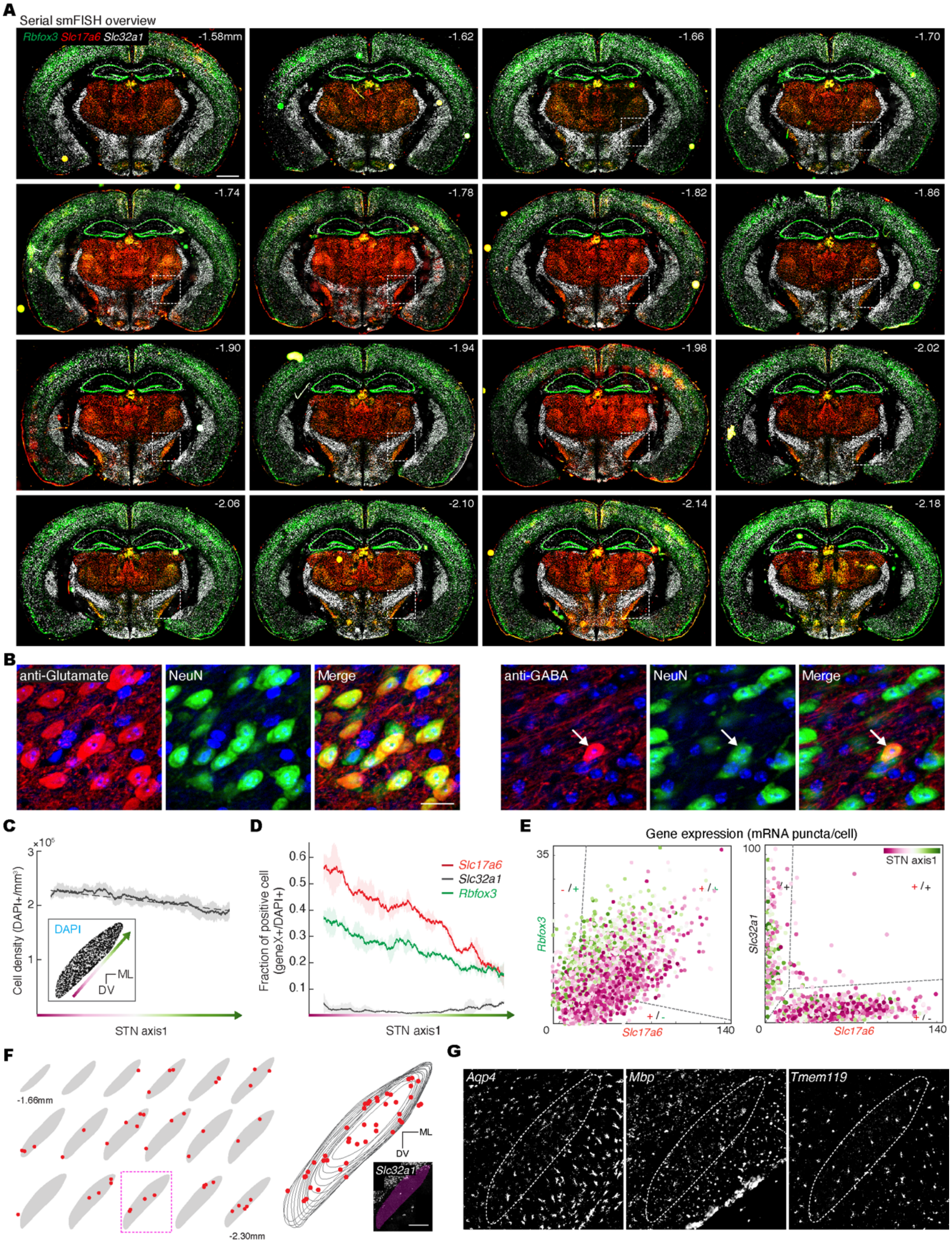


**Figure S7. Cellular composition of the STN by serial-smFISH, Related to Figure 4. (A)** Serial smFISH images of coronal sections containing the STN with *Rbfox3*, *Slc17a6*, and *Slc32a1* markers used to identify neurons, glutamatergic neurons, and GABAergic neurons, respectively (top). Enlarged view of the STN indicated by dashed boxes, with the distance from the bregma indicated at the corner. Scale bar, 1 mm (top); 250 μm (bottom). **(B)** Immunofluorescence image of STN cells showing co-labeling of anti-glutamate with NeuN (top) and anti-GABA with NeuN (bottom). Scale bar, 20 μm. **(C)** Average cell density (n = 10) along STN axis1, measured by the number of DAPI labeled cells (inset). **(D)** Quantification of cells expressing *Rbfox3*, *Slc17a6*, and *Slc32a* along STN axis1 (n = 2). **(E)** Co-expression plots of *Rbfox3* with *Slc17a6* (left) and *Slc32a1* with *Slc17a6* (right) showing co-expression of *Rbfox3* with *Slc17a6,* and distinct populations of Slc17a6- and Slc32a1-expressing cells. Data are represented as means with s.e.m. **(F)** Locations of GABAergic cells labeled by *Slc32a1* are shown on serial coronal STN sections ranging from −1.66 mm to −2.30 mm from bregma at 40-μm intervals (left). GABAergic cells are superimposed onto serial coronal STN boundaries (right). Scale bar, 250 μm. **(G)** Representative images of smFISH with *Aqp4, Mbp,* and *Tmem119* show three glial cell types: glia-astrocytes, oligodendrocytes, and microglia. Scale bar, 250 μm.

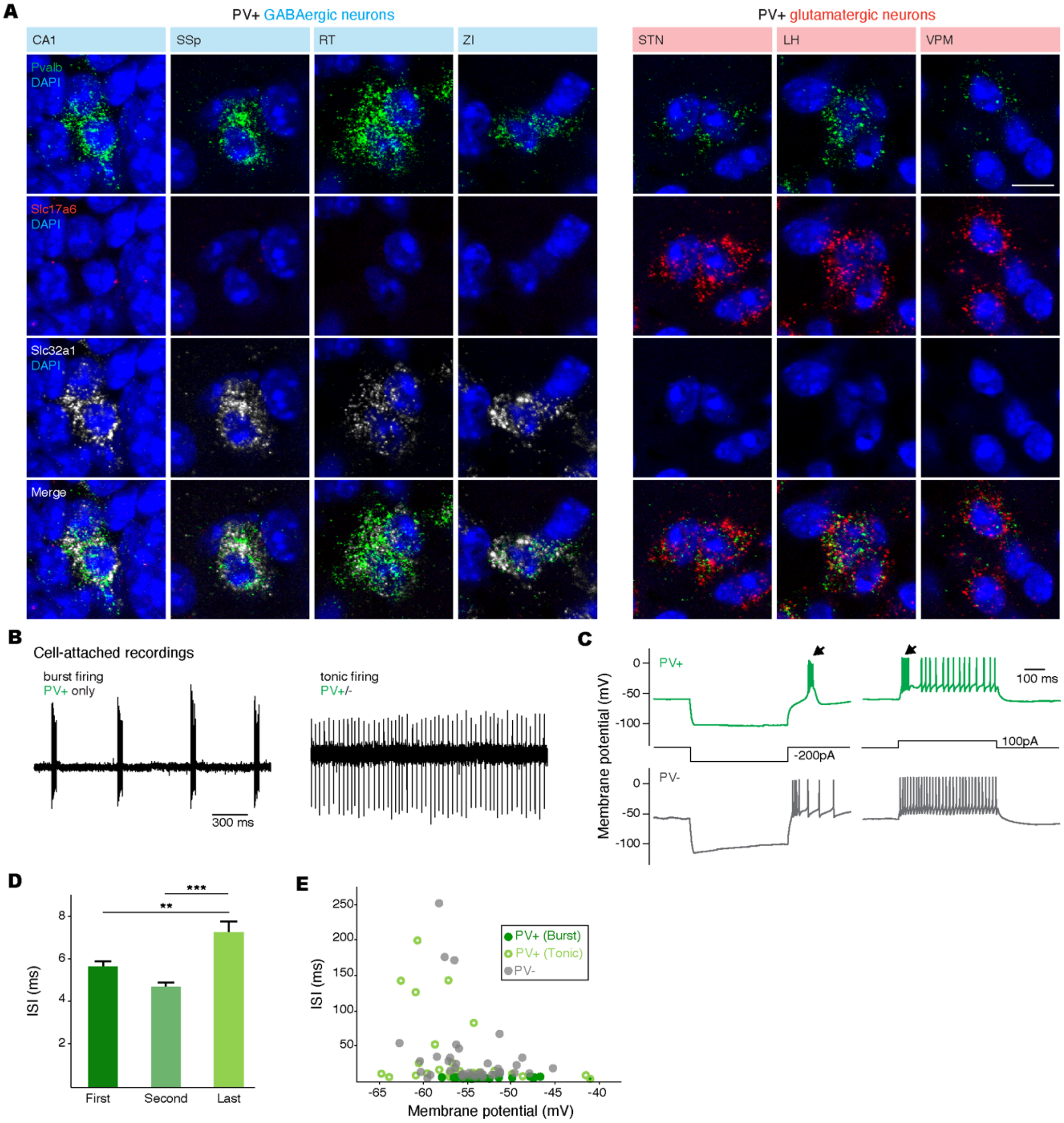


**Figure S8. Electrophysiological properties of STN PV+ neurons, Related to Figure 5. (A)** smFISH image of cells across the brain demonstrating GABAergic PV+ neurons in CA1, SSp, RT, and ZI co-expressing *Slc32a1* (left) and glutamatergic PV+ neurons in the STN, LH, and VPM co-expressing *Slc17a6* (right). Scale bar, 10 μm. **(B)** Representative burst (top) and tonic (bottom) firing recorded in cell-attached configurations. The tonic firing pattern was obtained from a STN PV− neuron, and the burst firing was obtained from a STN PV+ neuron. **(C)** Representative firing patterns in PV+ (top) and PV− (bottom) STN neurons when a hyperpolarization (left) or depolarization current (right) was injected. Arrows indicate burst firing. (**D**) ISI between the first two (first), 2nd and 3rd spikes (second), and last two spikes (last) during burst firing (n = 12). ** $p < 0.01$, *** $p < 0.001$, ANOVA followed by Turkey test. (**E**) Relationship between firing rate (ISI) and membrane potential in burst PV+ and tonic PV+/- neurons.

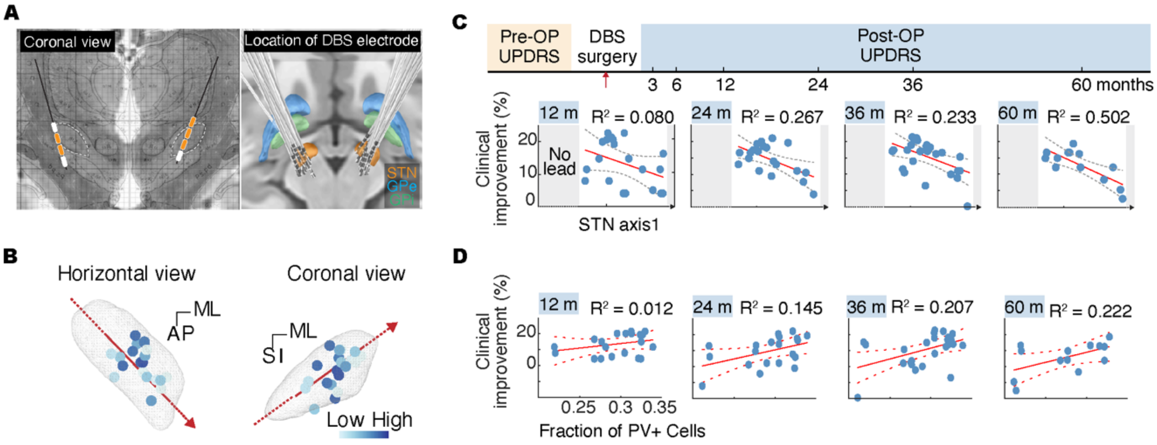


**Figure S9. Electrode position-dependent therapeutical effects in clinical DBS trials, Related to Main Text. (A)** Example DBS electrode position plotted on the coronal view of the human brain atlas of Schaltenbrand and Wahren (left). The yellow box denotes the activated lead. 3D visualization of superimposed electrode positions and trajectories from 26 patients who had undergone bilateral DBS surgery (right). **(B)** Positions of stimulated leads (n = 26) after surgery are superimposed onto the left hemisphere STN and colored based on improvements in motor symptom scores. **(C)** Illustration of DBS surgery and pre-/post-DBS surgery Unified Parkinson's Disease Rating Scale (UPDRS) evaluation timeline (top). Relationship between the DBS lead position along STN axis1 and improvements in clinical motor symptoms, measured by changes in the UPDRS III total score at 12, 24, 36, and 60 months after DBS surgery (bottom). Data were fitted using linear regression (red; adjusted R2 = 0.080, 0.267, 0.233, and 0.502, p = 0.12, 0.021, 0.003, and 0.005) with a 95% confidence bound (dotted). **(D)** Relationship between improvements in clinical motor symptoms after DBS hypothetical STN PV+ cell ratio at given electrode location from cross-species analysis. Data were fitted using linear regression (red; adjusted $R^2$ = 0.012, 0.145, 0.207, and 0.222, p = 0.283, 0.055, 0.017, and 0.059) with a 95% confidence bound (dotted).

#### Table S1. GPe injection summary

| | Injection Vol. (mm3) | | | | | | | | | | | | | | | | Projection Vol. (mm3) | Injection Coordinate | | | |
|---|---|---|---|---|---|---|---|---|---|---|---|---|---|---|---|---|---|---|---|---|---|
| ID | *int | AAA | RT | SI | STR | CEAc | CEAl | CEAm | CP | VPL | VPM | PAL | FS | GPe | GPi | em | STN | A-P | M-L | V-D | |
| JK311-1_sfGFP | | | | | | | | | 0.126 | | | | | 0.029 | | | 0.014 | 0 | 2.13 | -3.5 | Valid Injection |
| JK311-1_tdTomato | | | | | 0.002 | | | | 0.210 | | | 0.001 | | 0.105 | | | 0.029 | 0 | 1.87 | -3.5 | |
| JK311-2_sfGFP | | | | | 0.001 | | | | 0.187 | | | 0.000 | | 0.003 | | | 0.013 | 0 | 2.13 | -3.5 | |
| JK311-2_tdTomato | 0.000 | | | 0.000 | 0.003 | | | | 0.238 | | | 0.002 | | 0.019 | | | 0.024 | 0 | 1.87 | -3.5 | |
| JK311-3_sfGFP | | | | | 0.003 | | | | 0.201 | | | 0.002 | | 0.037 | | | 0.019 | 0 | 2.13 | -3.5 | |
| JK311-3_tdTomato | 0.001 | | | | 0.004 | | | | 0.148 | | | 0.002 | | 0.098 | | | 0.028 | 0 | 1.87 | -3.5 | |
| JK333-2_sfGFP | 0.002 | | | | | | | | 0.000 | | | 0.001 | | 0.040 | 0.000 | | 0.029 | -0.05 | 2 | -3.5 | |
| JK333-2_tdTomato | 0.060 | | 0.008 | | | | | | | | | 0.004 | | 0.058 | | 0.000 | 0.032 | -0.1 | 1.8 | -3.5 | |
| JK333-3_sfGFP | 0.001 | | | | | | | | 0.010 | | | | | 0.057 | | | 0.026 | -0.22 | 2.13 | -3.5 | |
| JK333-3_tdTomato | 0.063 | | 0.019 | | | | | | | 0.000 | | 0.006 | | 0.052 | | 0.000 | 0.029 | -0.22 | 1.87 | -3.5 | |
| JK335-1_sfGFP | 0.006 | | | | | | | | 0.014 | | | 0.001 | | 0.107 | 0.005 | | 0.030 | -0.22 | 1.88 | -3.43 | |
| JK338-1_sfGFP | 0.000 | | | | | | | | 0.007 | | | 0.001 | | 0.068 | 0.000 | | 0.029 | -0.22 | 1.88 | -3.68 | |
| JK338-1_tdTomato | 0.018 | | 0.000 | | | | | | | | | 0.003 | | 0.038 | 0.000 | | 0.010 | -0.22 | 1.63 | -3.68 | |
| JK352-1_sfGFP | 0.062 | | 0.020 | | | | | | | 0.002 | | 0.004 | | 0.038 | 0.001 | 0.002 | 0.013 | -0.22 | 1.63 | 3.93 | |
| JK352-2_sfGFP | | | | | 0.005 | | | | 0.159 | | | 0.002 | 0.000 | 0.053 | | | 0.018 | -0.22 | 1.88 | -3.93 | |
| JK352-2_tdTomato | 0.000 | | | | 0.000 | | | | 0.009 | | | 0.001 | | 0.075 | | | 0.034 | -0.22 | 1.63 | -3.68 | |
| JK352-5_sfGFP | | | | | 0.002 | | | | 0.085 | | | 0.002 | | 0.095 | | | 0.026 | -0.26 | 1.88 | -3.43 | |
| JK352-5_tdTomato | | | | | | | | | 0.013 | | | | | 0.063 | | | 0.036 | -0.34 | 2.13 | -3.5 | |
| JK356-5_sfGFP | 0.006 | | | | | | | | 0.008 | | | | | 0.063 | | | 0.022 | -0.34 | 1.87 | -3.5 | |
| JK356-6_sfGFP | | | | | 0.001 | | | | 0.009 | | | 0.005 | | 0.092 | 0.000 | | 0.031 | -0.34 | 2.15 | -3.5 | |
| JK358-1_sfGFP | 0.071 | | 0.036 | | | | | | | 0.000 | | 0.016 | | 0.026 | | 0.002 | 0.014 | -0.34 | 2 | -3.2 | |
| JK358-1_tdTomato | 0.011 | | | | | | | | 0.003 | | | 0.006 | | 0.088 | | | 0.031 | -0.34 | 2 | -4 | |
| JK358-5_sfGFP | 0.069 | | 0.018 | | | | | | | 0.001 | | 0.017 | | 0.049 | | 0.001 | 0.021 | -0.34 | 1.75 | -3.5 | |
| JK358-5_tdTomato | 0.066 | | 0.025 | | 0.000 | | | | | 0.000 | | 0.023 | | 0.052 | | 0.000 | 0.026 | -0.34 | 1.75 | -3.7 | |
| JK358-6_sfGFP | 0.000 | | | | 0.003 | | | | 0.012 | | | 0.003 | | 0.095 | | | 0.021 | -0.34 | 2.25 | -3.7 | |
| JK358-6_tdTomato | | | | 0.001 | 0.006 | | | | 0.091 | | | 0.007 | 0.000 | 0.041 | | | 0.025 | -0.34 | 1.8 | -4 | |
| JK359-1_sfGFP | 0.056 | | 0.070 | | | | | | | 0.040 | 0.000 | | | 0.013 | | 0.010 | 0.017 | -0.34 | 1.8 | -4 | |
| JK359-2_sfGFP | 0.007 | | | | | | | | 0.006 | | | 0.009 | | 0.121 | 0.005 | | 0.034 | -0.46 | 2.38 | -3.5 | |
| JK359-2_tdTomato | 0.065 | | 0.029 | | | | | | | 0.008 | | 0.017 | | 0.033 | 0.004 | 0.006 | 0.025 | -0.46 | 2.3 | -3.8 | |
| JK359-5_sfGFP | | | | | | | | | 0.106 | | | | | 0.036 | | | 0.028 | -0.46 | 2.3 | -3.8 | |
| JK359-6_sfGFP | 0.039 | | 0.004 | | | | | | | | | 0.009 | | 0.029 | 0.014 | | 0.011 | -0.46 | 2 | -3.8 | |
| JK359-6_tdTomato | 0.001 | | | 0.000 | 0.001 | | 0.000 | 0.000 | 0.008 | | | 0.003 | | 0.087 | 0.009 | | 0.031 | -0.46 | 2.12 | -3.25 | |
| JK363-1_sfGFP | 0.008 | | | 0.000 | | | | | | | | 0.031 | | 0.086 | 0.018 | | 0.021 | -0.46 | 2.12 | -3.75 | |
| JK363-2_sfGFP | 0.009 | | | | | | | | | | | 0.032 | | 0.031 | 0.029 | | 0.006 | -0.46 | 2.38 | -3.75 | |
| JK363-5_sfGFP | 0.034 | | 0.003 | | | | | | 0.002 | | | | | 0.045 | | | 0.012 | -0.58 | 2.26 | -3.1 | |
| JK363-5_tdTomato | 0.086 | | 0.065 | | 0.002 | | | | 0.002 | 0.014 | | 0.003 | | 0.025 | | 0.005 | 0.008 | -0.58 | 2 | -3.1 | |
| JK363-6_tdTomato | 0.003 | | | | | | | | 0.007 | | | | | 0.052 | | | 0.014 | -0.58 | 2.26 | -3.1 | |
| JK375-1_sfGFP | 0.014 | | | | | | | | 0.196 | | | | | 0.090 | | | 0.018 | -0.58 | 2.38 | -3.1 | |
| JK375-1_tdTomato | 0.014 | | | | | | | | 0.027 | | | 0.000 | | 0.086 | 0.000 | | 0.026 | -0.58 | 2.63 | -3.1 | |
| JK375-2_sfGFP | | | | | | | | | 0.141 | | | | | 0.071 | | | 0.029 | -0.58 | 2.38 | -3.35 | |
| JK375-2_tdTomato | | | | | 0.000 | | | | 0.188 | | | | | 0.040 | | | 0.019 | -0.58 | 2.38 | -3.35 | |
| JK375-5_tdTomato | 0.020 | | | | 0.000 | | | | 0.018 | | | 0.002 | | 0.103 | 0.011 | | 0.032 | -0.58 | 2.38 | -3.6 | |
| JK376-6_sfGFP | 0.033 | | 0.003 | | | | | | | | | 0.012 | | 0.030 | | | 0.003 | -0.7 | 2.38 | -3.5 | |
| JK416-1_sfGFP | 0.176 | | 0.124 | | | | | | 0.017 | 0.062 | 0.003 | 0.000 | | 0.062 | 0.001 | 0.016 | 0.007 | -0.7 | 2.12 | -3.5 | |
| JK416-1_tdTomato | 0.053 | | 0.011 | | | | | | 0.000 | | | | | 0.011 | | | 0.001 | -0.82 | 2.38 | -3 | |
| JK416-2_sfGFP | 0.024 | | | | 0.000 | | | | 0.054 | | | 0.000 | | 0.093 | 0.001 | | 0.018 | -0.82 | 2.62 | -3 | |
| JK416-2_tdTomato | 0.006 | | | 0.000 | 0.000 | | | | 0.006 | | | 0.002 | | 0.084 | 0.010 | | 0.025 | -0.82 | 2.38 | -3.5 | |
| JK453-2_sfGFP | | | | | | | | | 0.065 | | | | | 0.081 | | | 0.014 | -0.82 | 2.62 | -3.5 | |
| JK453-5_sfGFP | 0.039 | | 0.001 | | | | | | | | | 0.000 | | 0.029 | 0.002 | | 0.003 | -0.82 | 2.64 | -3.5 | |
| JK453-5_tdTomato | | | | | 0.008 | | 0.000 | 0.000 | 0.039 | | | 0.000 | | 0.030 | | | 0.001 | -0.82 | 2.4 | -3.5 | |
| JK453-6_sfGFP | 0.036 | | 0.002 | | | | | | 0.003 | | | 0.000 | | 0.051 | 0.000 | | 0.018 | -0.82 | 2.64 | -3.5 | |
| JK453-6_tdTomato | | | | | 0.000 | | | | 0.024 | | | | | 0.040 | | | 0.011 | -0.82 | 2.4 | -3.5 | |
| JK460-2_tdTomato | 0.015 | | | | | | | | | | | 0.000 | | 0.018 | 0.004 | | 0.004 | -0.82 | 2.64 | -3.5 | |
| JK460-3_tdTomato | 0.000 | | | 0.000 | 0.002 | | | 0.000 | 0.001 | | | 0.003 | | 0.039 | 0.005 | | 0.009 | -0.82 | 2.3 | -3.5 | |
| JK461-2_tdTomato | 0.013 | | | | 0.000 | | | | 0.000 | | | 0.002 | | 0.042 | | | 0.016 | -0.82 | 2.3 | -3.8 | |
| JK338-3_sfGFP | 0.001 | | | 0.025 | 0.008 | | 0.012 | 0.080 | 0.001 | | | 0.017 | | 0.036 | 0.008 | | 0.015 | | | | Ectopic injection |
| JK359-5_tdTomato | | | | | | | | | 0.301 | | | | | 0.004 | | | 0.020 | | | | |
| JK363-1_tdTomato | | 0.012 | | 0.005 | 0.038 | 0.007 | 0.006 | 0.060 | 0.047 | | | 0.010 | 0.013 | 0.028 | | | 0.026 | | | | |
| JK363-2_tdTomato | | 0.001 | | 0.005 | 0.016 | | 0.003 | 0.074 | 0.010 | | | 0.013 | 0.002 | 0.021 | 0.000 | | 0.027 | | | | |
| JK335-1_tdTomato | 0.003 | | | | | | | | | | | 0.000 | | 0.011 | | | 0.000 | | | | Weak STN projection / GPe injection |
| JK338-3_tdTomato | 0.020 | | | | | | | | | | | 0.006 | | 0.021 | 0.001 | | 0.000 | | | | |
| JK352-1_tdTomato | 0.000 | | 0.004 | | | | | | | | | 0.001 | | 0.000 | | | 0.000 | | | | |
| JK359-1_tdTomato | | | | | | | | | | | | | | 0.000 | | | 0.000 | | | | |
| JK363-6_sfGFP | 0.021 | | 0.012 | | | | | | | | | 0.000 | | 0.005 | | | 0.001 | | | | |
| JK376-6_tdTomato | 0.000 | | | | | | | | | | | 0.001 | | 0.009 | | | 0.000 | | | | |
| JK406-1_sfGFP | 0.011 | | 0.031 | | | | | | | 0.004 | | 0.005 | | 0.000 | | 0.005 | 0.000 | | | | |
| JK406-1_tdTomato | 0.015 | | 0.032 | | | | | | | 0.010 | | 0.002 | | 0.001 | | 0.004 | 0.003 | | | | |
| JK406-3_sfGFP | 0.045 | | 0.035 | | 0.001 | | | | 0.000 | 0.000 | | 0.016 | | 0.006 | | 0.000 | 0.000 | | | | |
| JK406-3_tdTomato | 0.043 | | 0.018 | | 0.003 | | | | 0.003 | | | 0.004 | | 0.006 | | 0.000 | 0.000 | | | | |
| JK453-2_tdTomato | | | | | | | | | 0.087 | | | | | 0.001 | | | 0.000 | | | | |
| JK460-2_sfGFP | 0.004 | | 0.018 | | | | | | | 0.013 | | | | 0.000 | | 0.004 | 0.000 | | | | |
| JK460-3_sfGFP | | | | | | | | | | | | | | 0.000 | | | 0.000 | | | | |
| JK460-4_sfGFP | | | | | 0.001 | | | | 0.000 | | | | | 0.003 | | | 0.000 | | | | |
| JK460-4_tdTomato | 0.008 | | | | | | | | | | | | | 0.002 | 0.000 | | 0.000 | | | | |
| JK461-2_sfGFP | | | | | | | | | | | | | | 0.000 | | | 0.000 | | | | |
| JK461-5_sfGFP | 0.033 | | 0.024 | 0.000 | 0.000 | | | | | | | 0.003 | | 0.014 | 0.011 | 0.000 | 0.001 | | | | |
| JK461-5_tdTomato | | | | | | | | | | | | | | 0.000 | | | 0.000 | | | | |

**Table S2.** Abbreviations

| Abbreviation | Name | CCF v3 id |
|---|---|---|
| **Mouse anatomy abbreviation** | | |
| CTX | Cerebral cortex | 688 |
| **Isocortex** | | |
| Isocortex | Isocortex | 315 |
| **Somatomotor cortex** | | |
| MO | Somatomotor areas | 500 |
| MOp | Primary motor area | 985 |
| MOs | Secondary motor area | 993 |
| SS | Somatosensory areas | 453 |
| SSp | Primary somatosensory area | 322 |
| SSp-n | Primary somatosensory area nose | 353 |
| SSp-bfd | Primary somatosensory area barrel field | 329 |
| SSp-ll | Primary somatosensory area lower limb | 337 |
| SSp-m | Primary somatosensory area mouth | 345 |
| SSp-ul | Primary somatosensory area upper limb | 369 |
| SSp-tr | Primary somatosensory area trunk | 361 |
| SSp-un | Primary somatosensory area unassigned | 182305689 |
| SSs | Supplemental somatosensory area | 378 |
| AI | Agranular insular area | 95 |
| AId | Agranular insular area dorsal part | 104 |
| AIp | Agranular insular area posterior part | 111 |
| AIv | Agranular insular area ventral part | 119 |
| **Association cortex** | | |
| GU | Gustatory areas | 1057 |
| VISC | Visceral area | 677 |
| AUD | Auditory areas | 247 |
| AUDd | Dorsal auditory area | 1011 |
| AUDp | Primary auditory area | 1002 |
| AUDpo | Posterior auditory area | 1027 |
| AUDv | Ventral auditory area | 1018 |
| VIS | Visual areas | 669 |
| RSP | Retrosplenial area | 254 |
| RSPagl | Retrosplenial area lateral agranular part | 894 |
| RSPd | Retrosplenial area dorsal part | 879 |
| RSPv | Retrosplenial area ventral part | 886 |
| PTLp | Posterior parietal association areas | 22 |
| TEa | Temporal association areas | 541 |
| PERI | Perirhinal area | 922 |
| ECT | Ectorhinal area | 895 |
| **Limbic cortex** | | |
| FRP | Frontal pole cerebral cortex | 184 |
| ACA | Anterior cingulate area | 31 |
| ACAd | Anterior cingulate area dorsal part | 39 |
| ACAv | Anterior cingulate area ventral part | 48 |
| PL | Prelimbic area | 972 |
| ILA | Infralimbic area | 44 |
| ORB | Orbital area | 714 |
| ORBl | Orbital area lateral part | 723 |
| ORBm | Orbital area medial part | 731 |
| ORBv | Orbital area ventral part | 738 |
| **Olfactory areas** | | |
| PIR | Piriform area | 961 |
| **Cortical plate** | | |

| | | |
|---|---|---|
| CLA | Claustrum | 583 |
| LA | Lateral amygdalar nucleus | 131 |
| BLA | Basolateral amygdalar nucleus | 295 |
| BLAv | Basolateral amygdalar nucleus ventral part | 451 |
| BMA | Basomedial amygdalar nucleus | 319 |
| **Striatum** | | |
| STR | Striatum | 477 |
| STRd | Striatum dorsal region | 485 |
| CP | Caudoputamen | 672 |
| STRv | Striatum ventral region | 493 |
| ACB | Nucleus accumbens | 56 |
| FS | Fundus of striatum | 998 |
| OT | Olfactory tubercle | 754 |
| AAA | Anterior amygdalar area | 23 |
| BA | Bed nucleus of the accessory olfactory tract | 292 |
| CEA | Central amygdalar nucleus | 536 |
| CEAc | Central amygdalar nucleus capsular part | 544 |
| CEAl | Central amygdalar nucleus lateral part | 551 |
| CEAm | Central amygdalar nucleus medial part | 559 |
| IA | Intercalated amygdalar nucleus | 1105 |
| MEA | Medial amygdalar nucleus | 403 |
| **Pallidum** | | |
| PAL | Pallidum | 803 |
| PALd | Pallidum dorsal region | 818 |
| GPe | Globus pallidus external segment | 1022 |
| GPi | Globus pallidus internal segment | 1031 |
| PALv | Pallidum ventral region | 835 |
| SI | Substantia innominata | 342 |
| MA | Magnocellular nucleus | 298 |
| PALm | Pallidum medial region | 826 |
| PALc | Pallidum caudal region | 809 |
| BST | Bed nuclei of the stria terminalis | 351 |
| BSTa | Bed nuclei of the stria terminalis anterior division | 359 |
| BAC | Bed nucleus of the anterior commissure | 287 |
| **Thalamus** | | |
| TH | Thalamus | 549 |
| DORsm | Thalamus sensory-motor cortex related | 864 |
| VENT | Ventral group of the dorsal thalamus | 637 |
| VAL | Ventral anterior-lateral complex of the thalamus | 629 |
| VM | Ventral medial nucleus of the thalamus | 685 |
| VP | Ventral posterior complex of the thalamus | 709 |
| VPL | Ventral posterolateral nucleus of the thalamus | 718 |
| VPM | Ventral posteromedial nucleus of the thalamus | 733 |
| SPF | Subparafascicular nucleus | 406 |
| SPFm | Subparafascicular nucleus magnocellular part | 414 |
| SPFp | Subparafascicular nucleus parvicellular part | 422 |
| SPA | Subparafascicular area | 609 |
| CM | Central medial nucleus of the thalamus | 599 |
| PF | Parafascicular nucleus | 930 |
| RT | Reticular nucleus of the thalamus | 262 |
| MH | Medial habenula | 483 |
| LH | Lateral habenula | 186 |
| **Hypothalamus** | | |
| HY | Hypothalamus | 1097 |
| LZ | Hypothalamic lateral zone | 290 |

| | | |
|---|---|---|
| LHA | Lateral hypothalamic area | 194 |
| PST | Preparasubthalamic nucleus | 356 |
| PSTN | Parasubthalamic nucleus | 364 |
| STN | Subthalamic nucleus | 470 |
| ZI | Zona incerta | 797 |
| **Midbrain** | | |
| MB | Midbrain | 313 |
| SCs | Superior colliculus sensory related | 302 |
| SCop | Superior colliculus optic layer | 851 |
| SCsg | Superior colliculus superficial gray layer | 842 |
| SCzo | Superior colliculus zonal layer | 834 |
| IC | Inferior colliculus | 4 |
| ICc | Inferior colliculus central nucleus | 811 |
| ICd | Inferior colliculus dorsal nucleus | 820 |
| ICe | Inferior colliculus external nucleus | 828 |
| VTA | Ventral tegmental area | 749 |
| SCm | Superior colliculus motor related | 294 |
| SCdg | Superior colliculus motor related deep gray layer | 26 |
| SCdw | Superior colliculus motor related deep white layer | 42 |
| SCiw | Superior colliculus motor related intermediate white layer | 17 |
| SCig | Superior colliculus motor related intermediate gray layer | 10 |
| PAG | Periaqueductal gray | 795 |
| RN | Red nucleus | 214 |
| SNl | Substantia nigra lateral part | 615 |
| SNc | Substantia nigra compact part | 374 |
| PPN | Pedunculopontine nucleus | 1052 |
| DR | Dorsal nucleus raphe | 872 |
| **Pons** | | |
| P | Pons | 771 |
| DTN | Dorsal tegmental nucleus | 880 |
| LTN | Lateral tegmental nucleus | 283 |
| LC | Locus ceruleus | 147 |
| LDT | Laterodorsal tegmental nucleus | 162 |
| **Fiber tracts** | | |
| cc | corpus callosum | 776 |
| ec | external capsule | 579 |
| cst | corticospinal tract | 784 |
| int | internal capsule | 6 |
| cpd | cerebal peduncle | 924 |
| V3 | third ventricle | 129 |
| V4 | fourth ventricle | 145 |
| act | anterior commissure temporal limb | 908 |
| em | external medullary lamina of the thalamus | 1092 |
| **Others** | | |
| HB | Hindbrain | 1065 |
| MY | Medulla | 354 |
| BS | Brain stem | 343 |
| IB | Interbrain | 1129 |

**Human anatomy abbreviation**

| Abbreviation | Definition | Brainnetome location |
|---|---|---|
| **Motor Cortices** | | |
| Mo | Motor | |
| M1 | Primary motor | A4hf + A4ul + A4t + A4tl +A4ll |

| | | |
|---|---|---|
| PM+SM | Premotor + Supplementary motor | A6dl + A6m + A6vl + A6cdl + A6cvl |
| **Prefrontal Association Cortices** | | |
| pA | Prefrontal association | |
| SFG | Superior frontal gyrus | A8m + A8dl + A9l + A9m + A10m |
| MFG | Middle frontal gyrus | A9/46d + IFJ + A46 + A9/46v + A8vl + A10l |
| IFG | Inferior frontal gyrus | A44d + IFS + A45c + A45r + A44op + A44v |
| **Rest Association Cortices** | | |
| reA | Rest association | |
| S | Somatosensory | A1/2/3ll + A1/2/3ulhf + A1/2/3tonIa + A2 + A1/2/3tru |
| SPL | Superior parietal lobule | A7r + A7c + A5l + A7pc + A7ip |
| IPL | Inferior parietal lobule | A39c + A39rd + A40rd + A40c + A39rv + A40rv |
| Pcun | Precuneus | A7m + A5m + dmPOS + A31 |
| MVOcC | MedioVentral occipital cortices | cLinG + rCunG + cCunG + rLinG + vmPOS |
| LOcC | Lateral occipital cortices | mOccG + V5/MT$_{+}$ + OPC + iOccG + msOccG + lsOccG |
| STG | Superior temporal gyrus | A38m + A41/42 + TE1.0 and TE1.2 + A22c + A38l + A22r |
| MTG | Middle temporal gyrus | A21c + A21r + A37dl + aSTS |
| ITG | Inferior temporal gyrus | A20iv + A37elv + A20r + A20il + A37vl + A20cl + A20cv |
| FuG | Fusiform gyrus | A20rv + A37mv + A37lv |
| PhG | Parahippocampal gyrus | A35/36r + TL + TI + TH |
| pSTS | Posterior superior temporal sulcus | rpSTS + cpSTS |
| INS | Insular gyrus | G + vIa + dIa+ vId/vIg + dIg + dId |
| **Limbic Cortices** | | |
| LC | Limbic cortices | |
| OrG | Orbital gyrus | A14m + A12/47o + A11l + A11m + A13 + A12/47l |
| CG | Cingulate gyrus | A23d + A24rv + A32p + A23v + A24cd + A23c + A32sg |
| EC | Entorhinal cortex | A35/36c + A28/34 |
| LN | Limbic nuclei | |
| Hipp | Hippocampus | |
| Amyg | Amygdala | |
| NAc | Nucleus accumbens | |
| Hyp | Hypothalamus | |
| MB | Mammillary body | |
| **Basal Ganglia** | | |
| CN | Caudate nucleus | |
| Pu | Putamen | |
| GPe | External globus pallidus | |
| GPi | Internal globus pallidus | |
| VP | Ventral pallidum | |
| STN | Subthalamic nucleus | |
| SN | Substantia nigra | |
| **White Matter** | | |
| IC | Internal capsule | |
| ICa | IC anterior limb | |

| | |
|---|---|
| ICg | IC genu |
| ICp | IC posterior limb |
| AC | Anterior commissure |
| PC | Posterior commissure |
| CrC | Crus cerebri |
| **Others** | |
| Thal | Thalamus |
| SC | Superior colliculus |
| RN | Red nucleus |

**Terminology**

| Abbreviation | Name |
|---|---|
| AAV | Adeno-Associated Virus |
| AHP | Afterhyperpolarizing potential |
| AMBCA | Allen Mouse Brain Connectivity |
| AP | Anterior-posterior |
| BG | Basal ganglia |
| CSD | Constrained spherical deconvolution |
| CT | Computed tomography |
| DBS | Deep brain stimulation |
| DV | Dorsal-ventral |
| E/I | Excitatory and inhibitory |
| HP | Hyperdirect pathway |
| $HP_{GPe}$ | Hyperdirect pathway projection in the GPe |
| $HP_{GPe/STN}$ | Hyperdirect pathway collateral projection in the GPe and the STN |
| $HP_{INPUT}$ | Hyperdirect pathway input in the cortex |
| $HP_{STN}$ | Hyperdirect pathway projection in the STN |
| IP | Indirect pathway |
| $IP_{INPUT}$ | Indirect pathway input from the GPe |
| $IP_{STN}$ | Indirect pathway projection in the STN |
| IT | Intratelencephalic |
| MDS | Multidimensional scaling |
| ML | Medial-lateral |
| MRI | Magnetic resonance imaging |
| MSE | Mean squared error |
| PD | Parkinson's disease |
| PT | Pyramidal tract |
| rAAV | Recombinant Adeno-Associated Virus |
| rCCA | Robust canonical correlation analysis |
| SI | Superior-inferior |
| smFISH | Single-molecule *in situ* hybridization |
| T2*WI | T2*-wighted imaging |
| TDI | Track-density imaging |
| UPDRS | Unified Parkinson Disease Rating Scale |

**Table S3.** List of transgenic Cre-lines used for hyperdirect pathway

**Ctx – STN**

| **Mouse line** | **Description** | **N** |
|---|---|---|
| Chrna2-Cre_OE25 | Enriched in cortical layer 5, olfactory areas, hippocampus, lateral septal complex, pallidum, thalamus, hypothalamus, midbrain, hindbrain, and cerebellum | 13 |
| Efr3a-Cre_NO108 | Enriched in cortical layers 5 and 6a, cortical subplate, thalamus, hypothalamus, medulla, pons, and cerebellum, striatum and midbrain. | 15 |
| Emx1-IRES-Cre | Enriched in cortex and hippocampus | 7 |
| Gpr26-Cre_KO250 | Enriched in frontal cortex, layer 5 and 6b in other cortical regions, CA1 of hippocampus, subnuclei of the thalamus, and in cerebellar Purkinje cells. Sparse expression in many other areas of the brain, including hindbrain, colliculus and striatum. | 7 |
| Htr2a-Cre_KM207 | Scattered expression throughout the brain. Enriched in layers 5 and 6b of cortex. Expression in restricted populations in septal complexes, hypothalamus and cerebellum. | 8 |
| Npr3-IRES2-Cre | Cre expression is enriched in layer 5 of cortex and in restricted populations within hippocampal formation. | 5 |
| Rbp4-Cre_KL100 | Enriched in cortical layer 5 and the dentate gyrus. | 46 |
| Sim1-Cre_KJ18 | Enriched in restricted populations within layer 5 of cortex, striatum (amygdala), and hypothalamus. Sparse, scattered expression elsewhere in the brain. | 14 |
| Syt6-Cre_KI148 | Sparse, scattered expression in brain areas including the medulla, pons, and midbrain. Enriched in specific areas within thalamus, layer 6a cortex, and olfactory areas. | 11 |
| Wild-type | | 55 |

**Ctx – GPe – STN**

| **Mouse line** | **Description** | **N** |
|---|---|---|
| Chrna2-Cre_OE25 | Enriched in cortical layer 5, olfactory areas, hippocampus, lateral septal complex, pallidum, thalamus, hypothalamus, midbrain, hindbrain, and cerebellum | 12 |
| Efr3a-Cre_NO108 | Enriched in cortical layers 5 and 6a, cortical subplate, thalamus, hypothalamus, medulla, pons, and cerebellum, striatum and midbrain. | 15 |
| Emx1-IRES-Cre | Enriched in cortex and hippocampus | 7 |
| Gpr26-Cre_KO250 | Enriched in frontal cortex, layer 5 and 6b in other cortical regions, CA1 of hippocampus, subnuclei of the thalamus, and in cerebellar Purkinje cells. Sparse expression in many other areas of the brain, including hindbrain, colliculus and striatum. | 7 |
| Htr2a-Cre_KM207 | Scattered expression throughout the brain. Enriched in layers 5 and 6b of cortex. Expression in restricted populations in septal complexes, hypothalamus and cerebellum. | 8 |
| Npr3-IRES2-Cre | Cre expression is enriched in layer 5 of cortex and in restricted populations within hippocampal formation. | 5 |
| Rbp4-Cre_KL100 | Enriched in cortical layer 5 and the dentate gyrus. | 46 |
| Sim1-Cre_KJ18 | Enriched in restricted populations within layer 5 of cortex, striatum (amygdala), and hypothalamus. Sparse, scattered expression elsewhere in the brain. | 13 |
| Syt6-Cre_KI148 | Sparse, scattered expression in brain areas including the medulla, pons, and midbrain. Enriched in specific areas within thalamus, layer 6a cortex, and olfactory areas. | 8 |
| Wild-type | | 55 |

**Table S4.** 7T-MRI parameters

| **Image** | [1] T1 | [2] DW | [3] T2* | [4] T2* | [5] T2* | [6] T1 |
|---|---|---|---|---|---|---|
| Scan parameters | | | | | | |
| Plane | sagittal | horizontal | horizontal | coronal | sagittal | 3D |
| Strength (T) | 7 | 7 | 7 | 7 | 7 | 3 |
| Sequence | MPRAGE | EPI | GRE | GRE | GRE | GR/IR |
| TR (ms) | 4000 | 6000 | 750 | 750 | 750 | 1900 |
| TE (ms) | 2.77 | 83 | 21.6 | 21.6 | 21.6 | 2.93 |
| FA (degree) | 10 | 90 | 30 | 30 | 30 | 9 |
| BW (Hz/pixel) | 470 | 1562 | 30 | 30 | 30 | 170 |
| Matrix | 256×256 | 128×128 | 1024×864 | 1024×864 | 1024×928 | 224×256 |
| Slice | 1 | 45 | 80 | 88 | 38 | 176 |
| Th (mm) | 4 | 1.8 | 2 | 2 | 2 | 1 |
| NEX | 32 | 3 | 1 | 1 | 1 | 1 |
| **Tractography** | (1) 033009 | (2) 033609 | (3) 0430045 | (4) 0436045 | (5) 043009 | (6) 043609 |
| Tracking parameters except the default options | | | | | | |
| Cutoff | 0.3 | 0.3 | 0.4 | 0.4 | 0.4 | 0.4 |
| Angle (degree) | 30 | 36 | 30 | 36 | 30 | 36 |
| Step (mm) | 0.9 | 0.9 | 0.45 | 0.45 | 0.9 | 0.9 |

**Table S5.** Electrophysiological parameters of STN PV+/- neurons

| **Parameters** | PV-negative | PV-positive | *p* value |
|---|---|---|---|
| Spontaneous firing | 35% (7/20 neurons) | 50% (13/26 neurons) | |
| Resting potential of silent neurons (mV) | -58.4 ± 1.2 (n=13) | -63.0 ± 2.3 (n = 13) | 0.243 |
| Threshold (mV) | -40.0 ± 0.6 (n = 20) | -40.2 ± 0.8 (n=26) | 0.882 |
| Cells showing tonic firing | | | |
| Inter-spike interval (ms) | 128 ± 48 (n=6) | 199 ± 31 (n=9) | 0.224 |
| Baseline (mV) | -48.7 ± 1.1 (n=6) | -49.0 ± 1.7 (n=9) | 0.689 |
| Overshoot (mV) | 16.7 ± 1.9 (n=6) | 12.1 ± 2.0 (n=9) | 0.0879 |
| AHP (mV) | -57.6 ± 1.1 (n=6) | -57.4 ± 1.5 (n=9) | 0.955 |
| Half width at half maximum (ms) | 0.328 ± 0.019 (n=6) | 0.293 ± 0.021 (n=9) | 0.327 |
| Threshold (mV) | -39.5 ± 0.9 (n=6) | -40.4 ± 1.5 (n=9) | 0.456 |
| Input resistance (MΩ) | 353.0 ± 48.2 (n = 20) | 419.6 ± 35.7 (n = 26) | 0.127 |

**Table S6.** Firing properties of burst and tonic firing neurons

| | | PV+ (burst) | PV+ (tonic) | PV- | Statistical test | | |
|---|---|---|---|---|---|---|---|
| | | | | | PV+ (burst) vs. PV+(tonic) | PV+ (burst) vs. PV- | PV+ (tonic) vs PV- |
| Examples (n) | Spontaneous | 4 | 6 | 7 | | | |
| | After the removal of -200 pA current injection | 9 | 9 | 12 | | | |
| | Upon 100 pA current injection | 7 | 14 | 19 | | | |
| ISI during burst firing or 3-6 spikes for tonic firing (ms) | Spontaneous | 5.6 ± 0.4 | 124.8 ± 19.2 | 112.7 ± 30.2 | * 0.031 | * 0.046 | 0.939 |
| | After the removal of -200 pA current injection# | 5.4 ± 0.3 | 13.6 ± 2.5 | 19.0 ± 4.9 | 0.33 | * 0.042 | 0.575 |
| | Upon 100 pA current injection# | 5.4 ± 0.4 | 11.4 ± 0.7 | 16.2 ± 1.8 | 0.088 | *** $5.49\times10^{-4}$ | 0.063 |
| Pause duration after burst firing by 100 pA current injection (ms) | | 92.4 ± 10.8 | – § | – § | | | |
| Hyperpolarization at the beginning of pause (mV) | Spontaneous | -20.4 ± 3.9 | – §§ | – §§ | | | |
| | After the removal of -200 pA current injection | -20.8 ± 2.3 | -1.5 ± 0.3 §§§ | -2.3 ± 0.5 §§§ | *** 0 | *** 0 | 0.897 |
| Increase in membrane potential during burst firing or 3-6 spikes for tonic firing (mV) | Spontaneous | 4.99 ± 1.16 | 0.056 ± 0.052 | -0.009 ± 0.05 | *** $7.48\times10^{-5}$ | *** $4.8\times10^{-5}$ | 0.995 |
| | After the removal of -200 pA current injection# | 7.57 ±0.86 | 0.78 ± 0.22 | 0.99 ± 0.52 | *** $2.83\times10^{-7}$ | *** $5.89\times10^{-8}$ | 1 |
| | Upon 100 pA current injection# | 7.56 ± 1.15 | 2.85 ± 0.67 | 2.27 ± 0.22 | *** $9.23\times10^{-5}$ | *** $6.95\times10^{-6}$ | 0.723 |

§ There are no clear pause time.
§§ There are no clear pause time.
§§§ There are no clear pause time and hyperpolarization. Instead, the difference in membrane potential between the third spike and the last spike during 250 ms after the removal of -200 pA current injection was calculated.
### For tonic firing neurons, average ISI or increase in membrane potential was calculated by using first 3 and 6 spikes for 100 pA current injection and the removal of -200 pA current injection, respectively.
*p < 0.05, *** p < 0.001, one-way ANOVA followed by Turkey test.

**Movie S1.** Aligned and registered 3D reconstructed whole brain slices using optimized mesoscopic mapping pipeline.

**Movie S2.** High resolution imaging and reconstruction of cleared STN of PV-IRES-Cre::Ai6 mice.
